## Supplementary figures and tables for "Volume Electron Microscopy Reveals Unique Laminar Synaptic Characteristics in the Human Entorhinal Cortex"

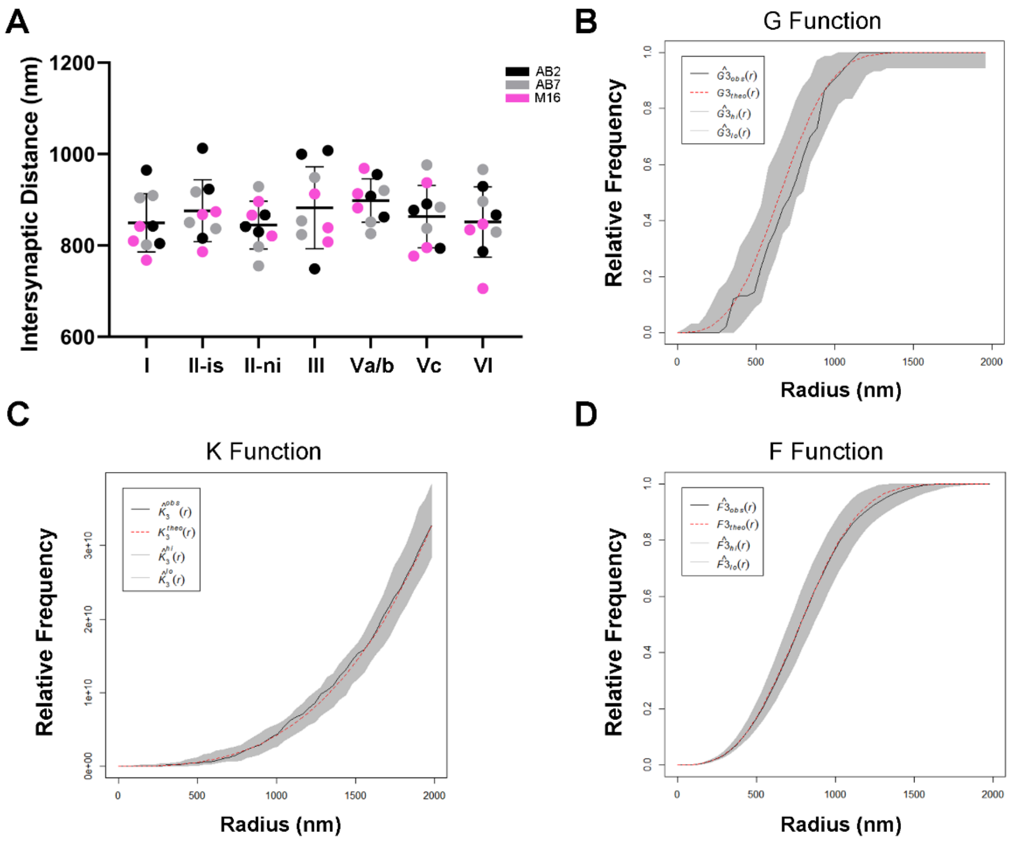


**Figure 1-figure supplement 1. Spatial Distribution analysis of synapses in the MEC.** (**A**) Mean intersynaptic distances (±SD) for each MEC layer. Each colored dot represents a stack of images from the analyzed cases AB2, AB7 and M16 (see **Supplementary File 1p** for details). No differences in the mean inter-synaptic distances were found between layers (KW; p>0.05). (**B–D**) Representative plots for G (**B**), K (**C**) and F (**D**) functions. Red dashed traces correspond to a theoretical homogeneous Poisson process for each function. The black continuous traces correspond to the experimentally observed function in the sample. The shaded areas represent the envelopes of values calculated from a set of 99 simulations.

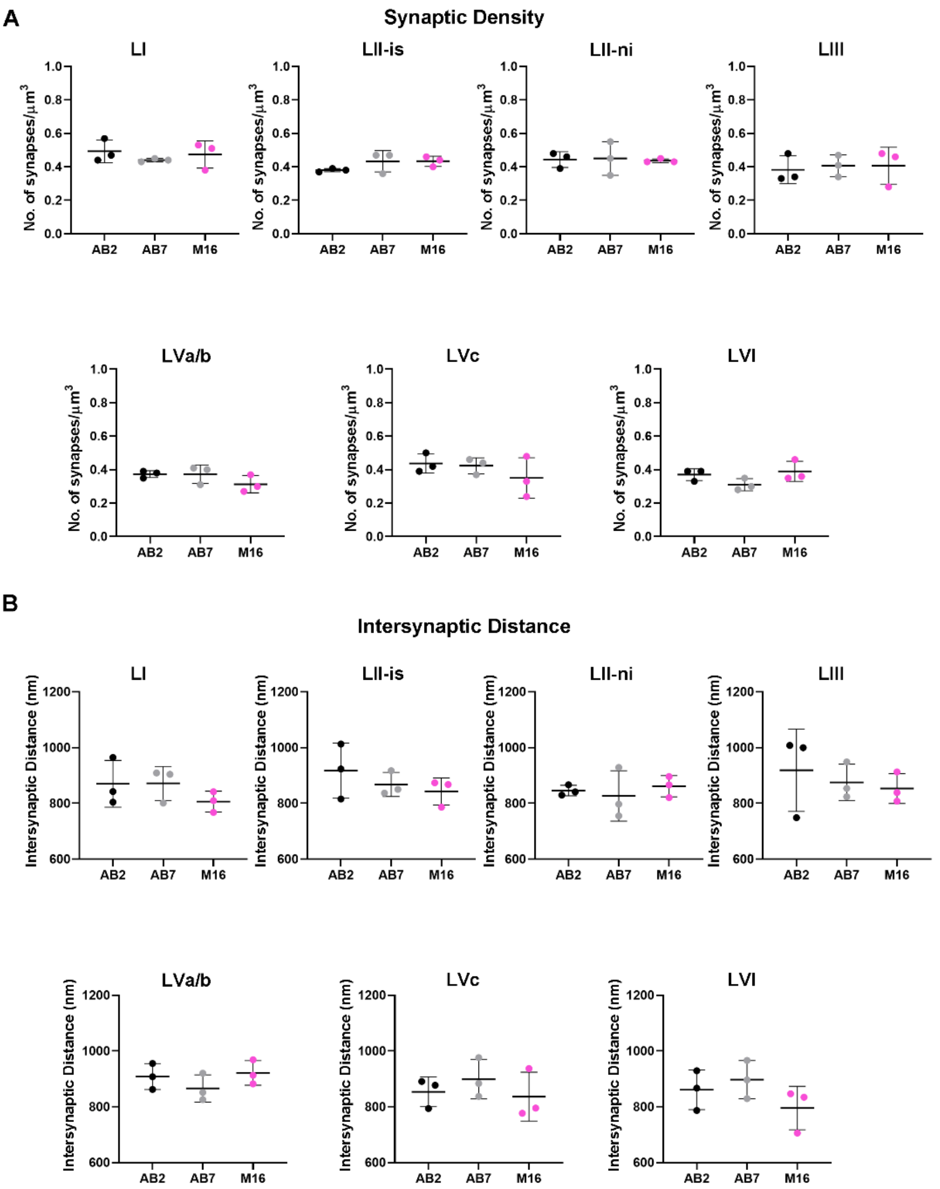


**Figure 1-figure supplement 2. Interindividual variability of synaptic density and intersynaptic distance in the MEC.** Separated plots per layer show the synaptic density (**A**) and intersynaptic distance (**B**) per case (mean±SD). No significant differences were found in any layer (KW, p>0.05). Each colored dot represents a stack of images from the analyzed cases AB2, AB7 and M16 (see **Supplementary File 1p** for details). P-values of comparisons are shown in **Supplementary File 3**.

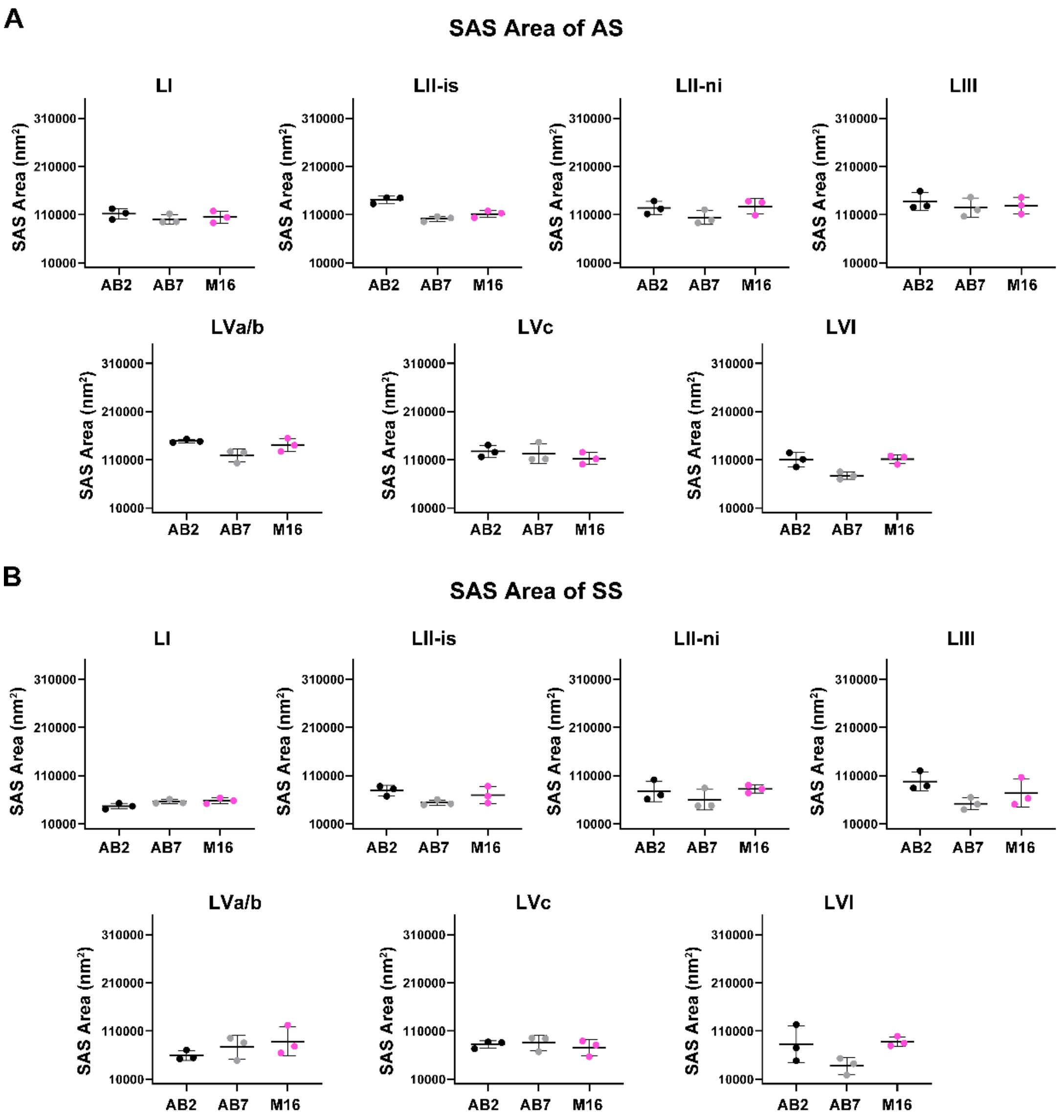


**Figure 3-figure supplement 1. Interindividual variability of SAS area of AS and SS in the MEC.** Separated plots per layer show the SAS area of AS (**A**) and SS (**B**) per case (mean±SD). In layer II-is, AB2 shows a larger SAS area of AS than AB7 (Dunn's test, $p<0.05$). No significant differences were found in the remaining comparisons (Dunn's test, $p>0.05$). Each colored dot represents a stack of images from the analyzed cases AB2, AB7 and M16 (see **Supplementary File 1p** for details). P-values of comparisons are shown in **Supplementary File 3.**

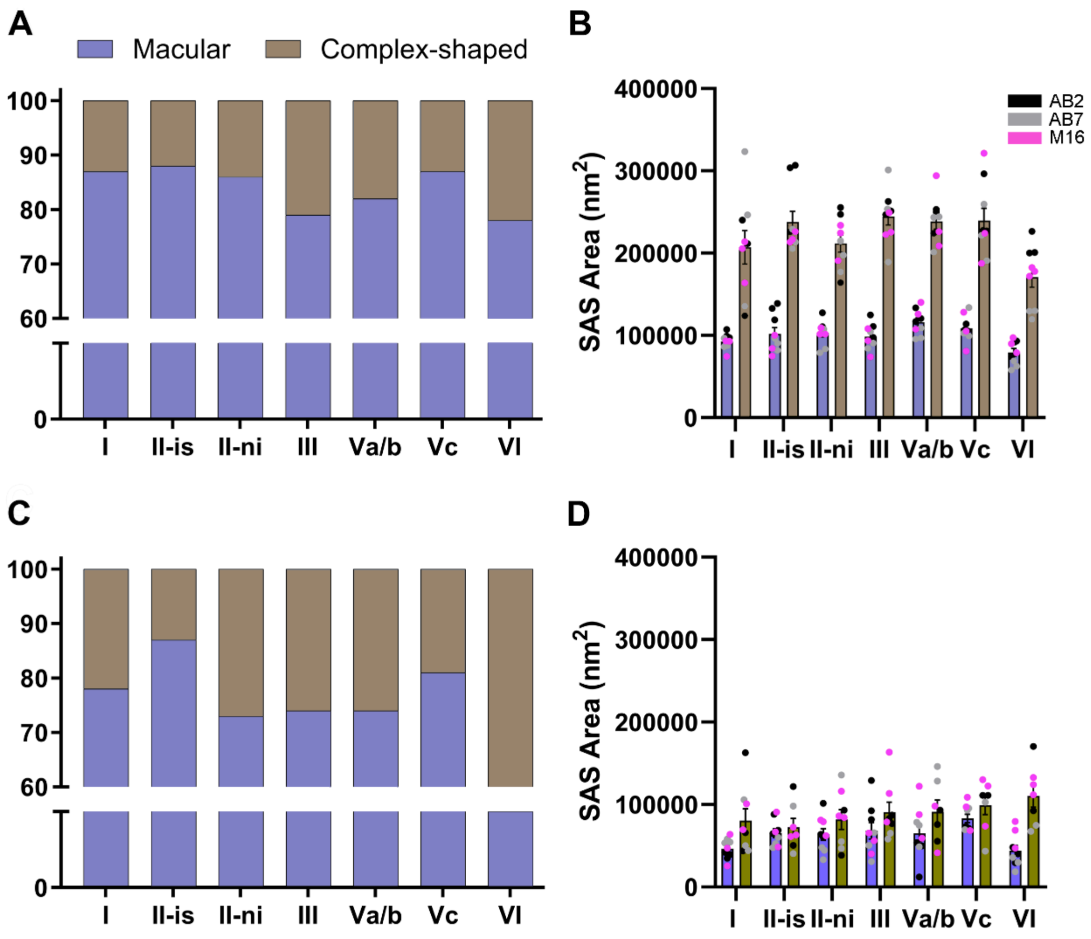


**Figure 4-figure supplement 1. Analysis of asymmetric (AS; A, B) and symmetric (SS; C, D) synapse shape in the MEC, per layer.** (A) Proportion of macular and complex-shaped AS (i.e., perforated, horseshoe and fragmented). Layer VI presented the lowest percentage of macular synapses (78%; $\chi^2$, p<0.0001). (B) Plot of the mean SAS area (±SE) of macular and complex-shaped AS. Complex-shaped AS were significantly larger than macular synapses (MW, p<0.05 in all layer comparisons). Each dot represents a stack of images from the analyzed cases AB2, AB7 and M16 (see **Supplementary File 1p** for details). (C) Proportion of macular and complex-shaped SS. Again, layer VI presented the lowest percentage of macular synapses (59%). (D) Plot of the mean SAS area (±SE) of macular and complex-shaped SS. Complex-shaped SS were larger than macular synapses in all layers; however, only layer I and layer VI presented significant differences (MW, p<0.05).

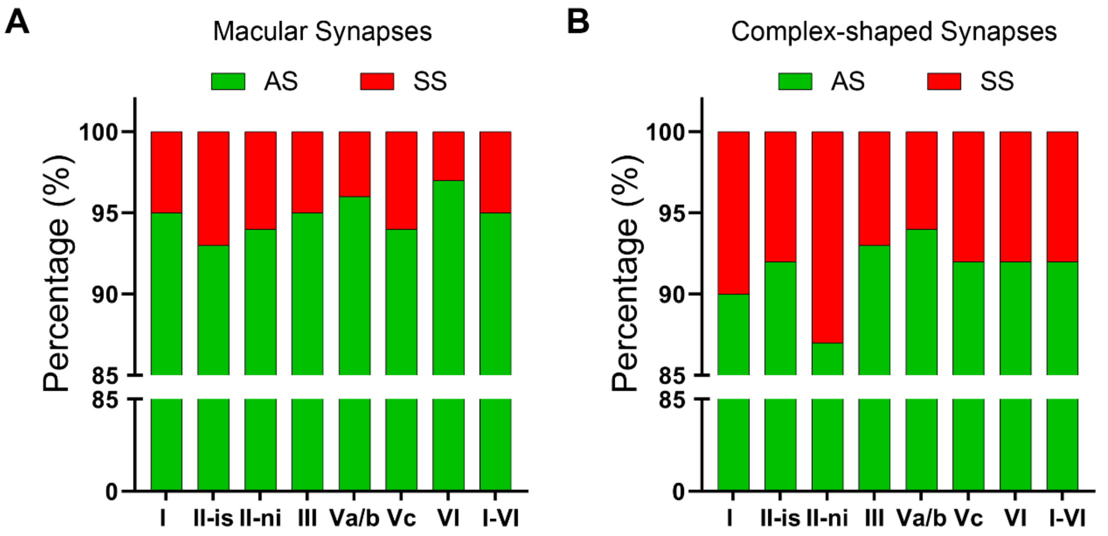


**Figure 4-figure supplement 2. Proportion of AS and SS in macular and complex-shaped synapses.** (**A**) Proportion of asymmetric (AS) and symmetric (SS) macular synapses. The overall AS:SS ratio of macular synapses was 95:5. No significant differences were found between layers ($\chi^2$, $p>0.0001$). (**B**) Proportion of AS and SS complex-shaped synapses. The overall ratio on complex-shaped synapses was 92:8. No significant differences were found between the layers ($\chi^2$, $p>0.0001$).

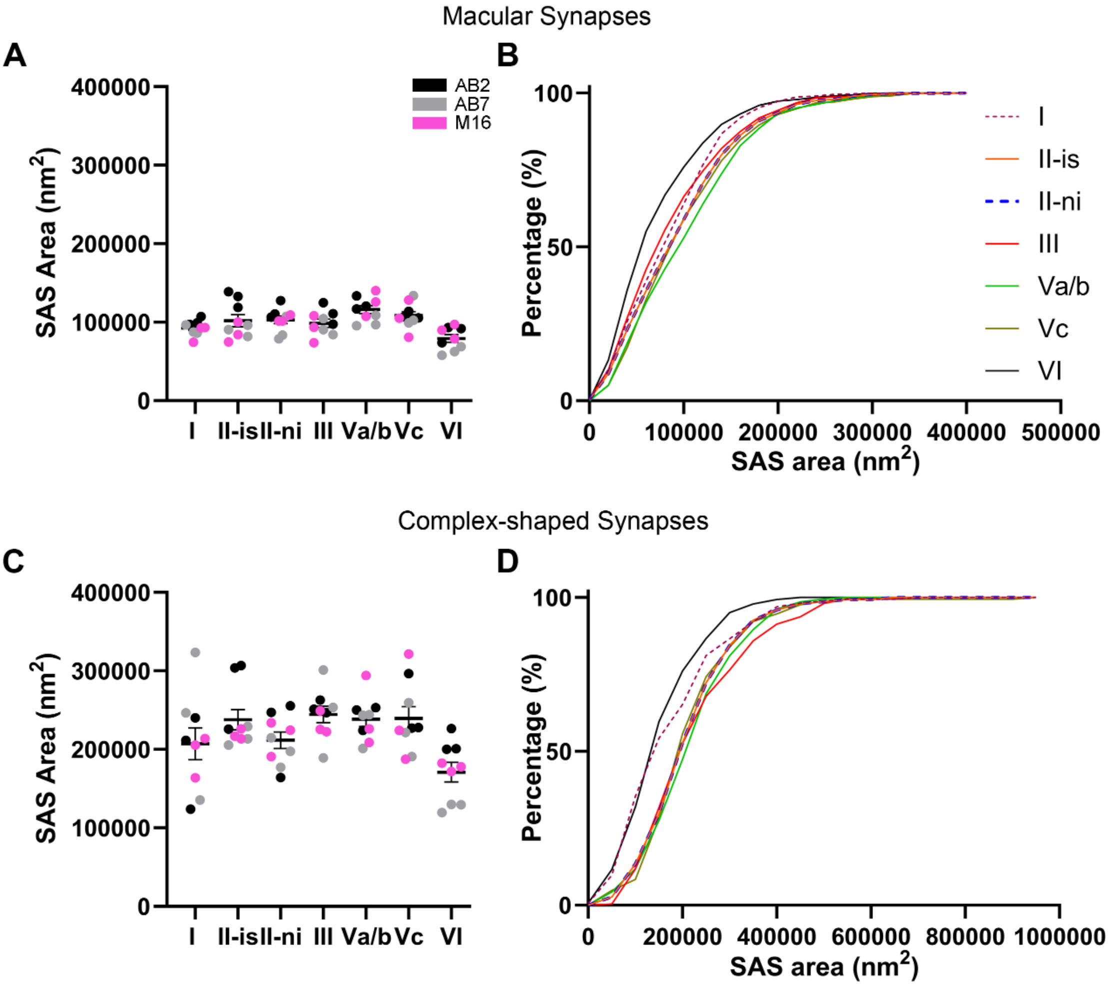


**Figure 4-figure supplement 3. Comparison of the SAS of macular (A, B) and complex-shaped (C, D) asymmetric synapses (AS) between MEC layers.** (**A**) Plots of the mean SAS area (±SE) of macular AS per layer. Layer VI had the smallest macular synapses of all layers (Dunn's test, $p<0.05$). Each dot represents a stack of images from the analyzed cases AB2, AB7 and M16 (see **Supplementary File 1p** for details). (**B**) Frequency distribution plot of SAS area per layer, for macular synapses, showing that smaller sizes were more frequent in layer VI (KS, $p<0.0001$). (**C**) Plots of the mean SAS area (±SE) of complex-shaped AS per layer. Again, layer VI had the smallest complex-shaped synapses of all layers (Dunn's test, $p<0.05$). (**D**) Frequency distribution plot of SAS area per layer, for complex synapses, showing that smaller sizes were more frequent in layer VI (KS, $p<0.0001$).

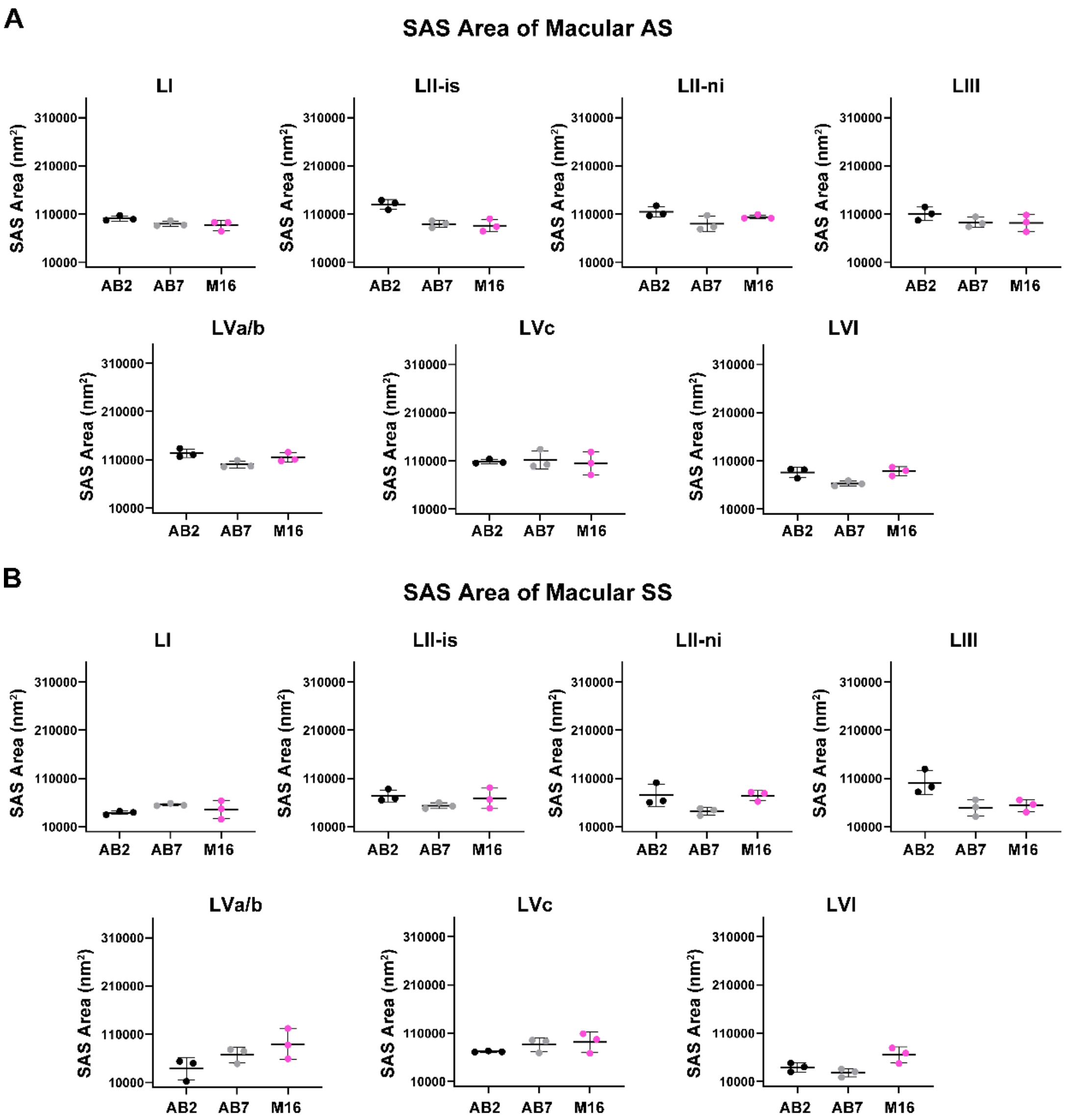


**Figure 4-figure supplement 4. Interindividual variability of SAS area of macular AS and SS in the MEC.** Separated plots per layer show the SAS area of macular AS (**A**) and SS (**B**) per case (mean±SD). No significant differences were found in any layer (KW, p>0.05). Each colored dot represents a stack of images from the analyzed cases AB2, AB7 and M16 (see **Supplementary File 1p** for details).  P-values of comparisons are shown in **Supplementary File 3**.

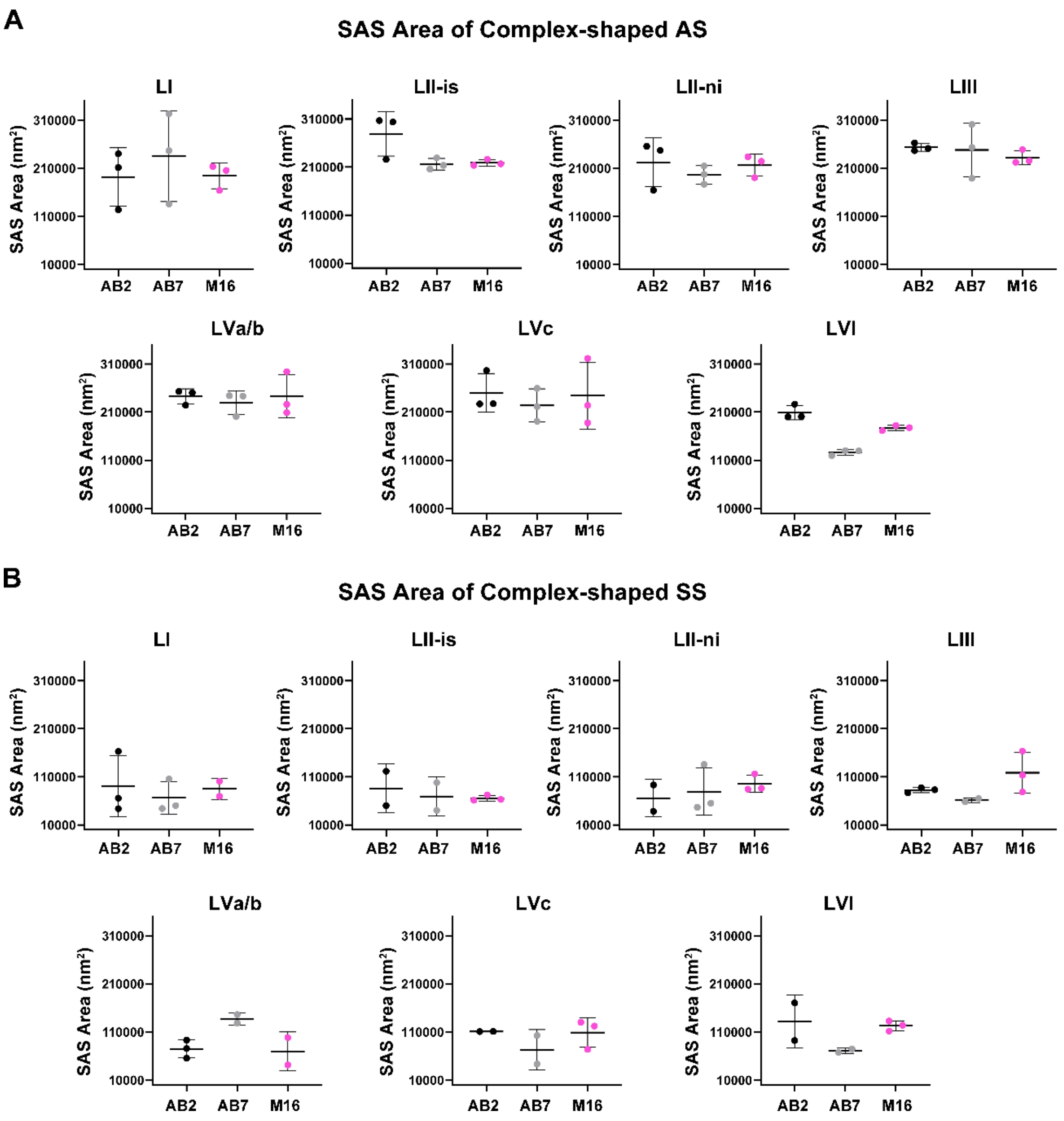


**Figure 4-figure supplement 5. Interindividual variability of SAS area of complex-shaped AS and SS in the MEC.** Separated plots per layer show the SAS area of complex-shaped AS (**A**) and SS (**B**) per case (mean±SD). In LVI, AB2 have larger SAS area of complex-shaped AS than AB7 (Dunn's test, $p<0.05$). No significant differences were found in the rest of the layers (Dunn's test, $p>0.05$). Each colored dot represents a stack of images from the analyzed cases AB2, AB7 and M16 (see **Supplementary File 1p** for details). P-values of comparisons are shown in **Supplementary File 3**.

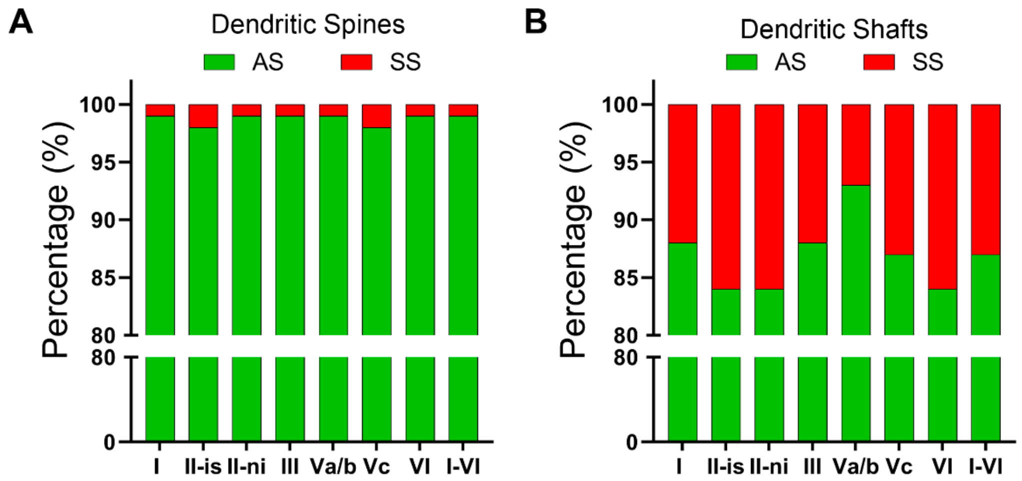


**Figure 6-figure supplement 1. Proportion of AS and SS in dendritic spines and shafts.** (**A**) Proportion of asymmetric (AS) and symmetric (SS) synapses on dendritic spines. The overall AS:SS ratio was 99:1. No significant differences were found between the layers ($\chi^2$, p>0.0001). (**B**) Proportion of AS and SS on dendritic shafts. The overall ratio was 87:13. No significant differences were found between the layers ($\chi^2$, p>0.0001).

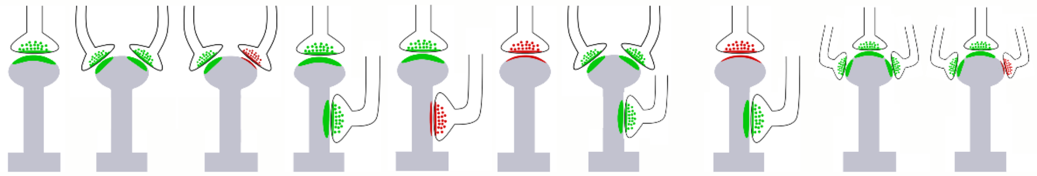

| | | | | | | | | | | |
|---|---|---|---|---|---|---|---|---|---|---|
| I<br>(672) | 96.43%<br>(649) | 1.49%<br>(10) | 0.15%<br>(1) | 0.59%<br>(4) | 0.45%<br>(3) | 0.74%<br>(5) | -<br>(0) | 0.15%<br>(1) | -<br>(0) | -<br>(0) |
| II-is<br>(737) | 95.25%<br>(702) | 1.36%<br>(10) | 1.09%<br>(8) | 0.95%<br>(7) | 0.27%<br>(2) | 0.95%<br>(7) | 0.14%<br>(1) | -<br>(0) | -<br>(0) | -<br>(0) |
| II-ni<br>(797) | 97.24%<br>(775) | 1.25%<br>(10) | 0.50%<br>(4) | 0.38%<br>(3) | 0.25%<br>(2) | 0.38%<br>(3) | -<br>(0) | -<br>(0) | -<br>(0) | -<br>(0) |
| III<br>(659) | 95.90%<br>(632) | 1.82%<br>(12) | 1.06%<br>(7) | 0.91%<br>(6) | -<br>(0) | 0.30%<br>(2) | -<br>(0) | -<br>(0) | -<br>(0) | -<br>(0) |
| Va/b<br>(567) | 96.12%<br>(545) | 1.23%<br>(7) | -<br>(0) | 1.94%<br>(11) | 0.35%<br>(2) | 0.18%<br>(1) | -<br>(0) | -<br>(0) | -<br>(0) | 0.18%<br>(1) |
| Vc<br>(772) | 94.30%<br>(728) | 1.94%<br>(15) | 1.81%<br>(14) | 1.17%<br>(9) | 0.13%<br>(1) | 0.52%<br>(4) | -<br>(0) | -<br>(0) | -<br>(0) | 0.13%<br>(1) |
| VI<br>(843) | 97.51%<br>(822) | 1.19%<br>(10) | 0.36%<br>(3) | 0.36%<br>(3) | 0.12%<br>(1) | 0.36%<br>(3) | -<br>(0) | -<br>(0) | 0.12%<br>(1) | -<br>(0) |
| **I-VI<br>(5048)** | **96.13%<br>(4853)** | **1.47%<br>(74)** | **0.73%<br>(37)** | **0.85%<br>(43)** | **0.22%<br>(11)** | **0.50%<br>(25)** | **0.02%<br>(1)** | **0.02%<br>(1)** | **0.02%<br>(1)** | **0.04%<br>(2)** |

**Figure 6-figure supplement 2. Schematic representation of the proportion of single and multi-synaptic spines in all MEC layers.** Data in parentheses refer to the absolute number of spines.

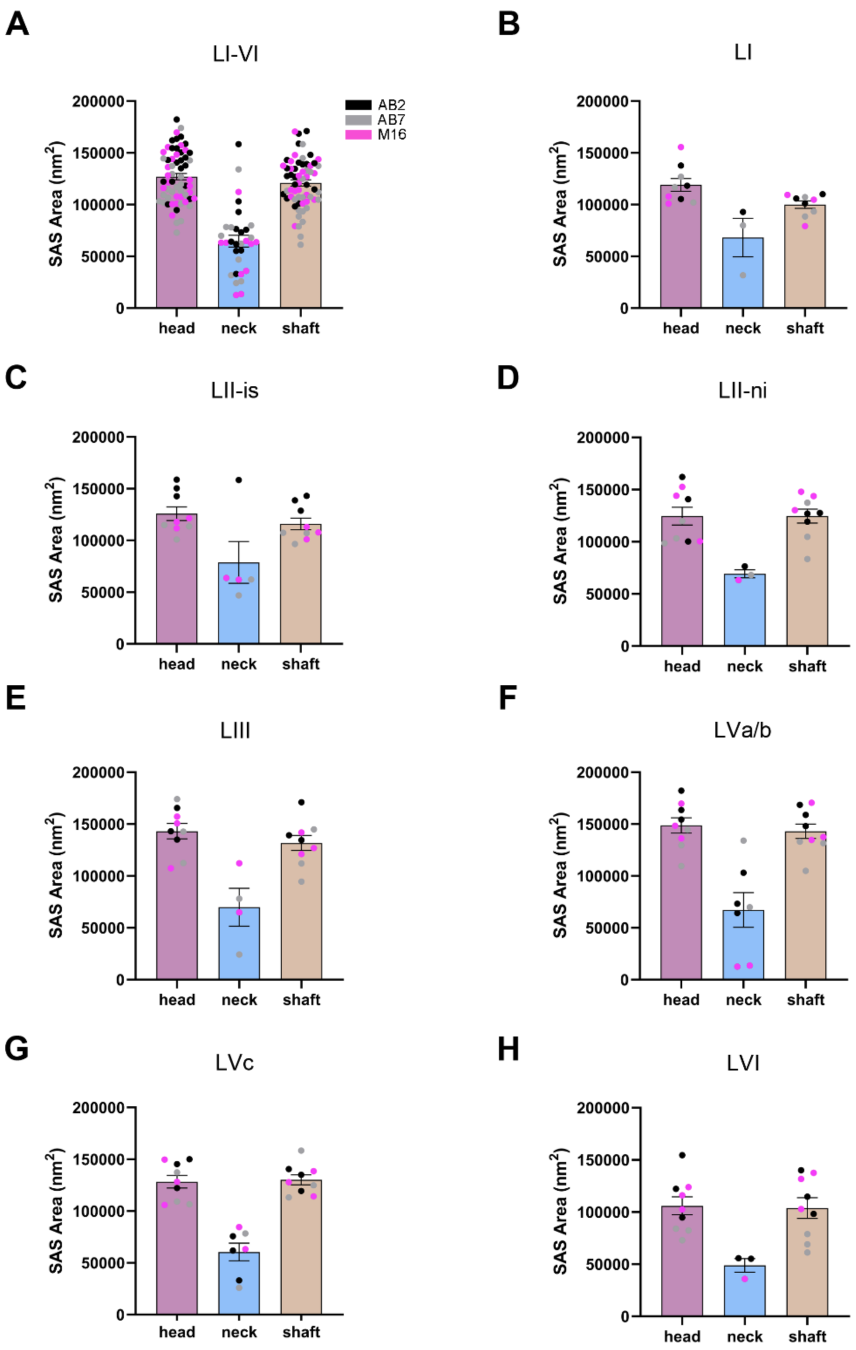


**Figure 6-figure supplement 3. Analysis of SAS of asymmetric synapses (AS) for each of the post-synaptic targets in the MEC.** (**A**) Plots of the mean SAS area (±SE) for AS established on spine heads, necks and dendritic shafts, considering all layers. Each dot represents a stack of images from the analyzed cases AB2, AB7 and M16 (see **Supplementary File 1p** for details). Synapses established on the spine neck were significantly smaller (Dunn's test, $p<0.001$). (**B–H**) Plots of the mean SAS area (±SE) for AS established on spine heads, necks and dendritic shafts of each MEC layer.

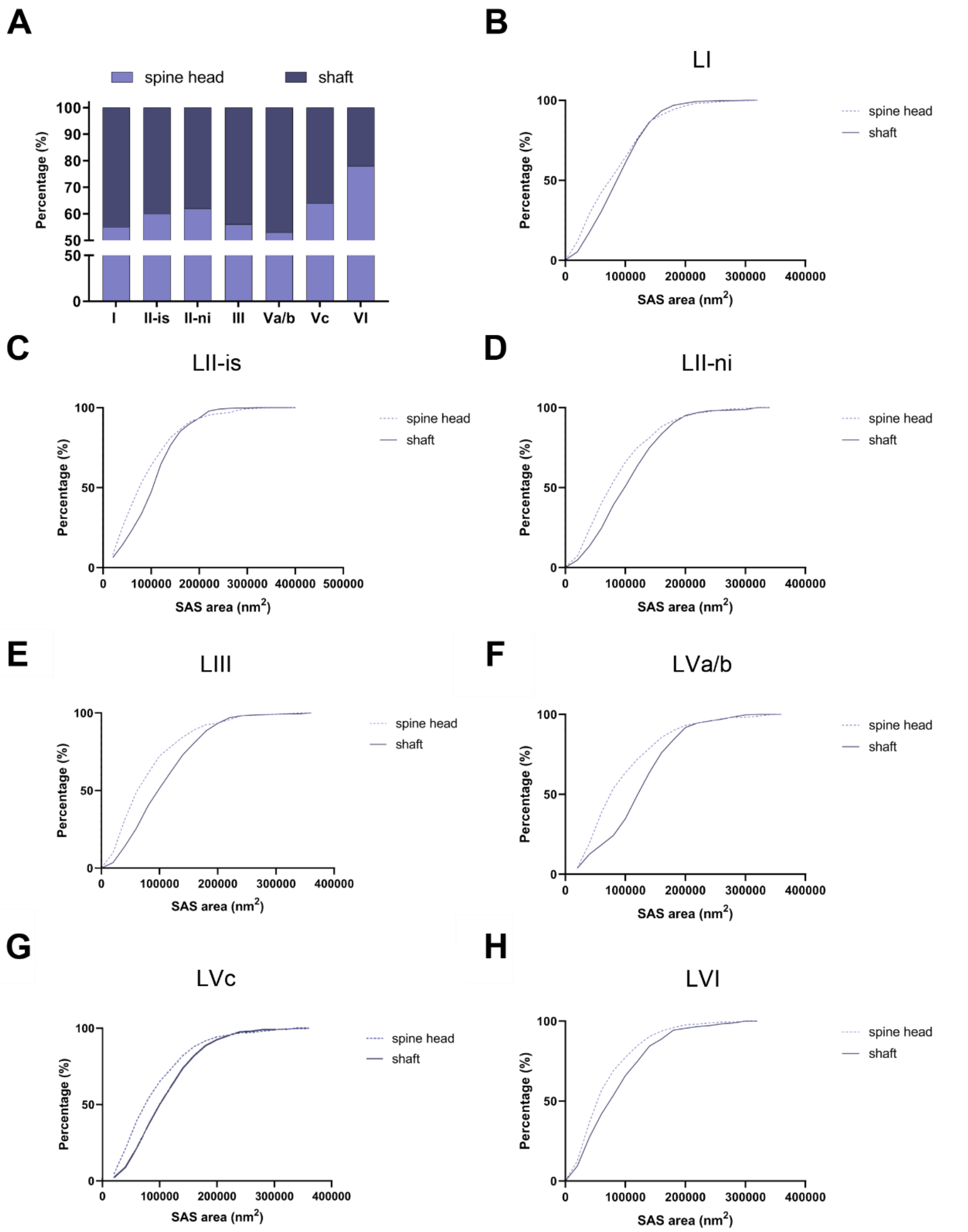


**Figure 7-figure supplement 1. Analysis of the distribution of macular AS on post-synaptic targets, per MEC layer. (A)** Distribution of macular AS on spine heads and dendritic shafts in each layer. Layer VI exhibited the highest proportion of macular AS established on spine heads (78%; $\chi^2$, p<0.0001). **(B–H)** Frequency distribution plots of the SAS of macular AS established on spine heads and dendritic shafts, per cortical layer.

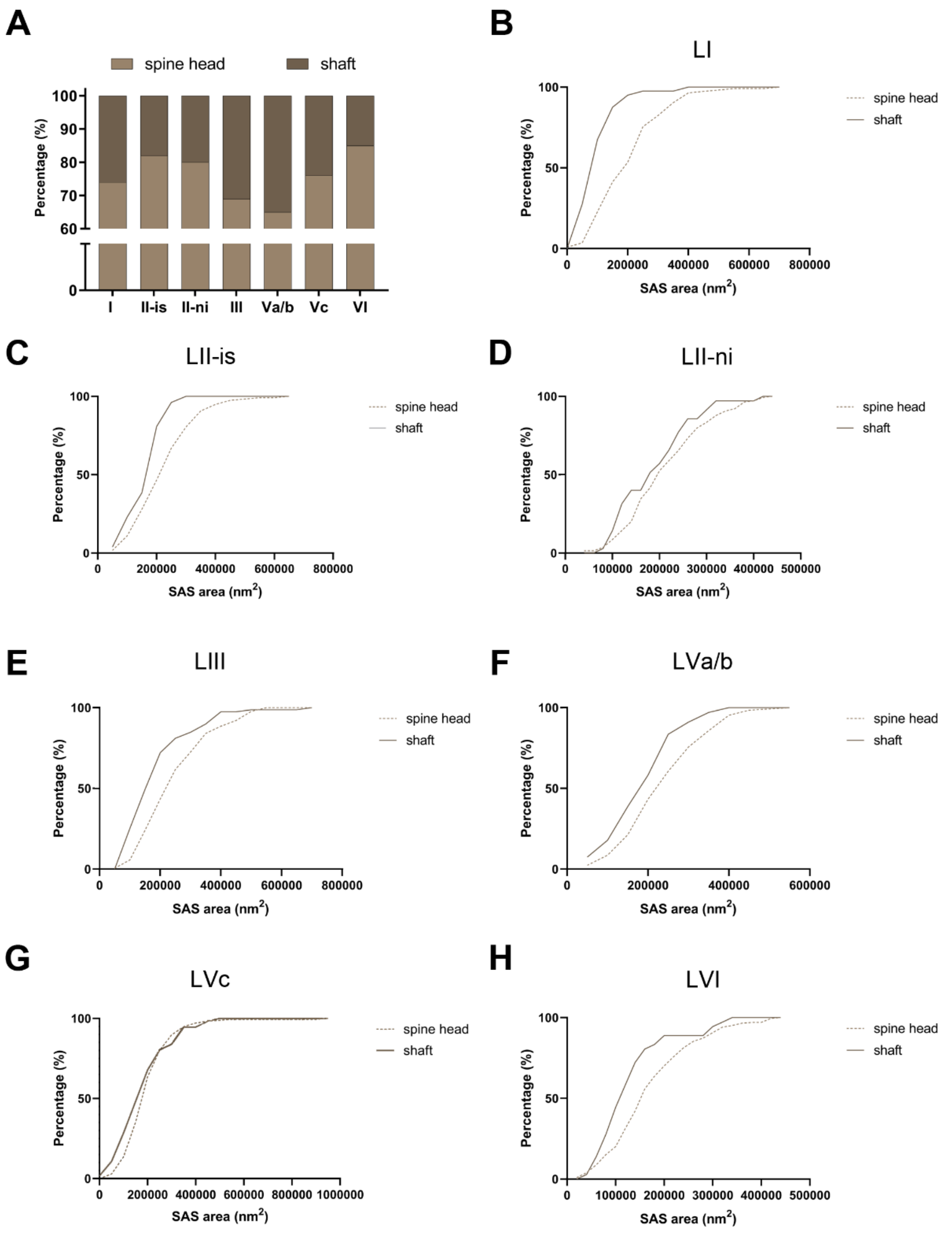


**Figure 7-figure supplement 2. Analysis of the distribution of complex-shaped AS on post-synaptic targets, per MEC layer. (A)** Distribution of complex-shaped AS on spine heads and dendritic shafts in each layer. Layer VI exhibited the highest proportion of complex-shaped AS established on spine heads (85%, $\chi^2$, p<0.0001). **(B–H)** Frequency distribution plots of the SAS of complex-shaped AS established on spine heads and dendritic shafts, per cortical layer.

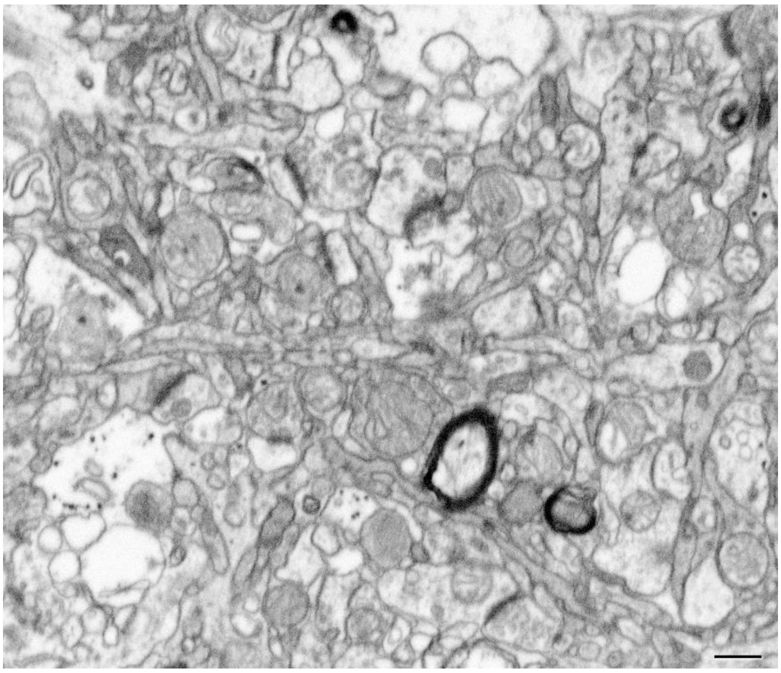

**Figure 10-figure supplement 1. Ultrastructure of the neuropil in layer I.** FIB/SEM image resolution in the xy plane was 5 nm/pixel. Resolution in the z-axis (section thickness) was 20 nm. Scale bar indicates 575 nm.

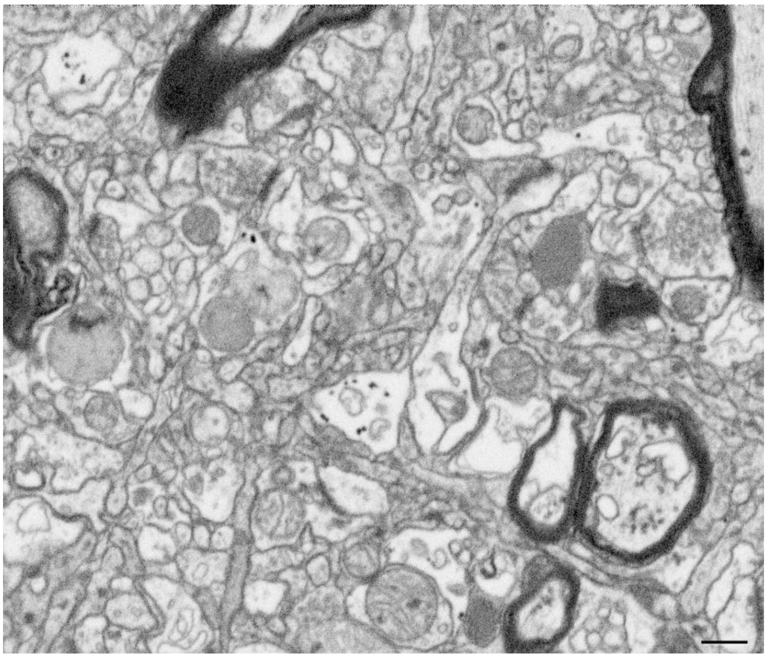

**Figure 10-figure supplement 2. Ultrastructure of the neuropil in layer II-is.** FIB/SEM image resolution in the xy plane was 5 nm/pixel. Resolution in the z-axis (section thickness) was 20 nm. Scale bar indicates 575 nm.

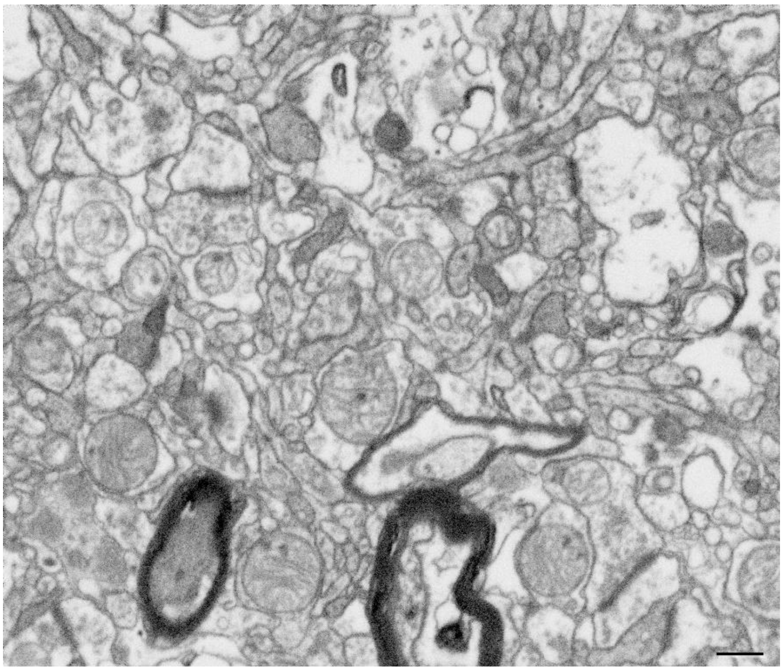

**Figure 10-figure supplement 3. Ultrastructure of the neuropil in layer II-ni.** FIB/SEM image resolution in the xy plane was 5 nm/pixel. Resolution in the z-axis (section thickness) was 20 nm. Scale bar indicates 575 nm.

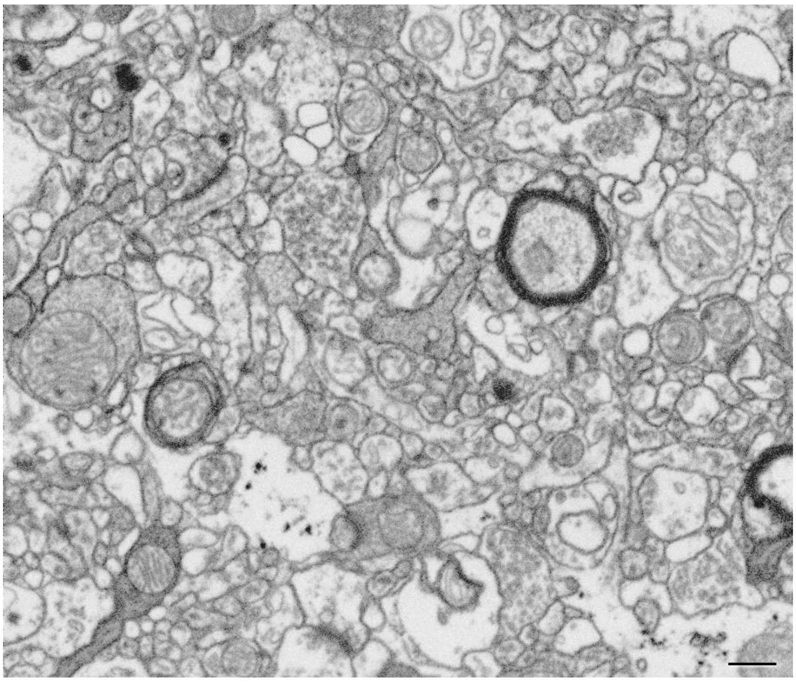

**Figure 10-figure supplement 4. Ultrastructure of the neuropil in layer III.** FIB/SEM image resolution in the xy plane was 5 nm/pixel. Resolution in the z-axis (section thickness) was 20 nm. Scale bar indicates 575 nm.

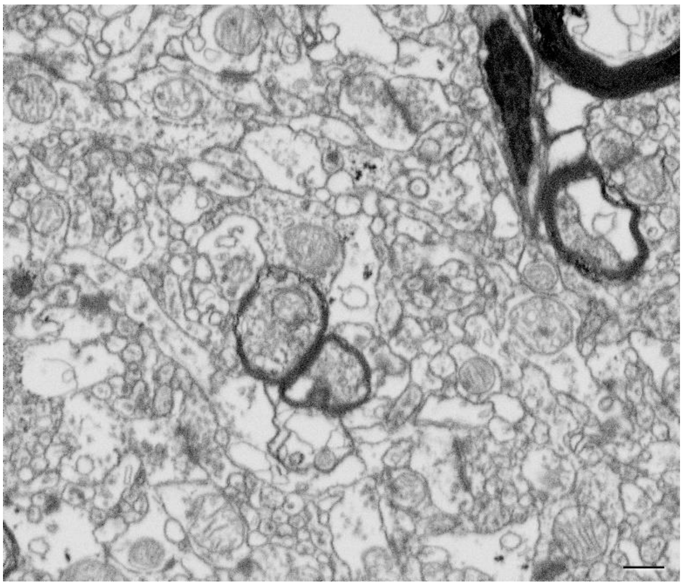

**Figure 10-figure supplement 5. Ultrastructure of the neuropil in layer Vc.** FIB/SEM image resolution in the xy plane was 5 nm/pixel. Resolution in the z-axis (section thickness) was 20 nm. Scale bar indicates 575 nm.

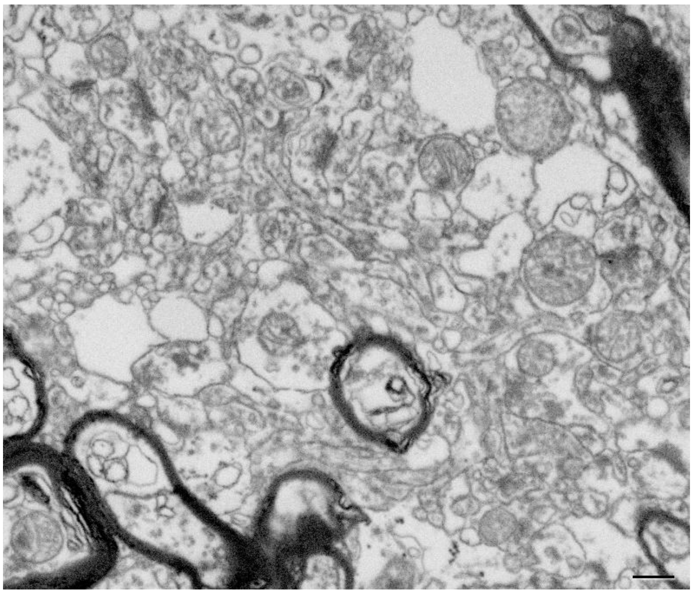

**Figure 10-figure supplement 6. Ultrastructure of the neuropil in layer VI.** FIB/SEM image resolution in the xy plane was 5 nm/pixel. Resolution in the z-axis (section thickness) was 20 nm. Scale bar indicates 575 nm.

**Legend to Figure 13-Video 1. Visualization of the reconstructed dendritic segment of figure 13, using EspINA software.** Please consider that some image quality has been lost during the video compression process.

| Layer | $V_c$ (%±SD) | $V_{bv}$ (%±SD) | $V_n$ (%±SD) |
|---|---|---|---|
| **I** | 1.40±0.43 | 2.70±0.62 | 96.01±0.25 |
| **II-is** | 9.20±0.66 | 2.69±1.45 | 88.11±2.03 |
| **II-ni** | 3.13±1.04 | 4.38±0.43 | 92.49±1.13 |
| **III** | 9.26±0.91 | 5.53±1.74 | 85.71±0.83 |
| **Va/b** | 8.97±1.05 | 3.57±0.45 | 87.46±1.07 |
| **Vc** | 6.80±1.25 | 2.86±0.11 | 90.34±1.33 |
| **VI** | 6.60±0.75 | 3.99±1.27 | 89.41±2.02 |

**Supplementary File 1a. Light microscopy data: volume fraction occupied by cortical elements in layers I, II-is, II-ni, III, Va/b, Vc and VI of the MEC.** Vc: Volume fraction occupied by cells bodies; Vbv: Volume fraction occupied by blood vessels; Vn: Volume fraction occupied by neuropil.

| Layer | Case | $V_c$ (%) | $V_{bv}$ (%) | $V_n$ (%) |
|---|---|---|---|---|
| I | **AB2** | 1.82 | 2.01 | 96.17 |
| | **AB7** | 0.97 | 3.20 | 95.83 |
| | **M16** | 1.42 | 2.88 | 95.70 |
| II-is | **AB2** | 9.75 | 4.33 | 85.92 |
| | **AB7** | 9.40 | 2.11 | 88.49 |
| | **M16** | 8.46 | 1.62 | 89.92 |
| II-ni | **AB2** | 3.68 | 4.82 | 91.50 |
| | **AB7** | 3.78 | 3.97 | 92.25 |
| | **M16** | 1.93 | 4.35 | 93.72 |
| III | **AB2** | 9.70 | 4.70 | 85.60 |
| | **AB7** | 8.21 | 7.48 | 84.31 |
| | **M16** | 9.87 | 4.31 | 85.82 |
| Va/b | **AB2** | 7.92 | 3.40 | 88.68 |
| | **AB7** | 10.03 | 3.24 | 86.73 |
| | **M16** | 8.95 | 4.08 | 86.97 |
| Vc | **AB2** | 8.01 | 2.98 | 89.01 |
| | **AB7** | 6.87 | 2.79 | 90.36 |
| | **M16** | 5.51 | 2.83 | 91.66 |
| VI | **AB2** | 7.36 | 5.35 | 87.28 |
| | **AB7** | 5.86 | 2.85 | 91.30 |
| | **M16** | 6.58 | 3.76 | 89.67 |

**Supplementary File 1b. Light microscopy data: volume fraction occupied by cortical elements in layers I, II-is, II-ni, III, Va/b, Vc and VI of the MEC, for individual cases.** Vc: Volume fraction occupied by cells bodies; Vbv: Volume fraction occupied by blood vessels; Vn: Volume fraction occupied by neuropil.

| Layer | Case | No. of stack of images | Total volume of Neuropil ($\mu m^3$) | No. of images per stack (range) | Total No. of Images |
|---|---|---|---|---|---|
| **I** | AB2, AB7, M16 | 9 | 4195 | 271-308 | 2667 |
| **II-is** | AB2, AB7, M16 | 9 | 4048 | 262-300 | 2574 |
| **II-ni** | AB2, AB7, M16 | 9 | 3984 | 244-320 | 2533 |
| **III** | AB2, AB7, M16 | 9 | 3990 | 254-293 | 2537 |
| **Va/b** | AB2, AB7, M16 | 9 | 4051 | 270-311 | 2576 |
| **Vc** | AB2, AB7, M16 | 9 | 4095 | 229-302 | 2557 |
| **VI** | AB2, AB7, M16 | 9 | 4113 | 271-319 | 2615 |
| **I-VI** | - | 63 | 28,476 | - | 18,059 |

**Supplementary File 1c. Summary of the stack details obtained from the multiple sampling from all layers of MEC.** 3 stacks of images were acquired in each MEC layer, per case (9 stacks of images in total per layer). The last row shows the sum of all the MEC layers from all cases.

| Layer | Case | No. of AS | No. of SS | No. all synapses | % AS (mean) | % SS (mean) | CFs volume (μm³) | No. AS /μm³ (mean±SD) | No. SS /μm³ (mean±SD) | No. all synapses/μm³ (mean±SD) | Area of SAS AS (nm²; mean ± SE) | Area of SAS SS (nm²; mean ± SE) | Intersynaptic distance (nm; mean±SD) |
|---|---|---|---|---|---|---|---|---|---|---|---|---|---|
| I | **AB2** | 422 | 31 | 453 | 93 | 7 | 825 (1,073) | 0.46±0.07 (0.39±0.07) | 0.03±0.01 (0.03±0.00) | 0.49±0.07 (0.42±0.06) | 111,057±6,462 (103,283±6,010) | 46,632±3,449 (43,368±3,208) | 871±84 (844±81) |
| | **AB7** | 424 | 23 | 447 | 94.9 | 5.1 | 905 (1,117) | 0.42±0.01 (0.38±0.02) | 0.02±001 (0.02±0.00) | 0.44±0.01 (0.40±0.02) | 111,785±5,563 (93,730±5,174) | 55,767±2,562 (51,864±2,383) | 872±61 (846±59) |
| | **M16** | 419 | 26 | 445 | 94.1 | 5.9 | 844 (1,074) | 0.45±0.08 (0.39±0.06) | 0.03±0.01 (0.02±0.01) | 0.47±0.08 (0.41±0.06) | 105,134±7,055 (97,774±6,561) | 57,701±3,480 (53,662±3,237) | 806±37 (782±36) |
| II-is | **AB2** | 423 | 37 | 460 | 91.9 | 8.1 | 1089 (1,122) | 0.35±0.02 (0.38±0.02) | 0.03±0.00 (0.03±0.01) | 0.38±0.01 (0.41±0.01) | 140,997±4,257 (131,127±3,959) | 79,196±6,205 (73,652±5,771) | 917±99 (890±96) |
| | **AB7** | 427 | 30 | 457 | 93.4 | 6.6 | 951 (974) | 0.40±0.06 (0.44±0.07) | 0.03±0.01 (0.03±0.01) | 0.43±0.06 (0.47±0.07) | 101,566±3,152 (94,456±2,932 | 53,705±3,259 (49,946±3,031) | 868±43 (842±42) |
| | **M16** | 402 | 33 | 435 | 92.5 | 7.5 | 909 (1,045) | 0.40±0.02 (0.38±0.02) | 0.03±0.01 (0.03±0.01) | 0.43±0.03 (0.42±0.04) | 112,125±4,434 (104,276±4,124) | 71,862±12,206 (66,832±11,351) | 843±49 (817±47) |
| II-ni | **AB2** | 428 | 26 | 454 | 94.4 | 5.6 | 927 (988) | 0.42±0.03 (0.43±0.04 | 0.02±0.01 (0.02±0.01) | 0.44±0.05 (0.46±0.05) | 124,176±7,895 (115,484±7,342) | 77,432±12,221 (72,012±11,366) | 846±19 (821±18) |
| | **AB7** | 483 | 26 | 509 | 95.2 | 4.8 | 1023 (1,083) | 0.42±0.09 (0.45±0.09) | 0.02±0.01 (0.03±0.01) | 0.45±0.10 (0.47±0.11) | 104,123±8,321 (96,834±7,739) | 59,852±12,299 (55,663±11,438) | 827±90 (802±88) |
| | **M16** | 379 | 45 | 424 | 89.5 | 10.5 | 870 (947) | 0.39±0.01 (0.40±0.02) | 0.04±0.01 (0.05±0.01) | 0.44±0.01 (0.45±0.01) | 127,715±9,094 (118,775±8,458) | 82,140±4,689 (76,391±4,360) | 861±38 (835±37) |
| III | **AB2** | 390 | 21 | 411 | 94.8 | 5.2 | 961 (995) | 0.36±0.08 (0.39±0.07) | 0.02±0.01 (0.02±0.01) | 0.38±0.08 (0.41±0.07) | 138,360±10 187 (128,675±9,474) | 97,730±11,236 (90,889±10,449) | 919±147 (891±143) |
| | **AB7** | 415 | 21 | 436 | 95.3 | 4.7 | 958 (999) | 0.39±0.06 (0.41±0.07) | 0.02±0.01 (0.02±0.01) | 0.41±0.06 (0.44±0.06) | 124,549±11,540 (115,830±10,732) | 51,655±7,341 (48,039±6,827) | 875±65 (849±63) |
| | **M16** | 411 | 32 | 443 | 92,3 | 7.7 | 982 (1,019) | 0.38±0.11 (0.40±0.11) | 0.03±0.01 (0.03±0.01) | 0.41±0.11 (0.43±0.11) | 128,981±10,056 (119,952±9,352) | 73,365±16,872 (68,229±15,691) | 853±54 (828±52) |

| | | | | | | | | | | | | | |
|---|---|---|---|---|---|---|---|---|---|---|---|---|---|
| Va/b | **AB2** | 418 | 18 | 436 | 95.8 | 4.2 | 1051 (1,083) | 0.36±0.03 (0.39±0.03) | 0.02±0.006 (0.02±0.006) | 0.37±0.02 (0.40±0.03) | 149,010±2,110 (138,579±1,963) | 59,037±5,827 (54,904±5,419) | 908±46 (881±45) |
| | **AB7** | 414 | 17 | 431 | 96 | 4 | 1034 (1,059) | 0.36±0.05 (0.39±0.05) | 0.01±0.006 (0.01±0.006) | 0.37±0.06 (0.40±0.06) | 118,640±7,720 (110,335±7,180) | 76,658±14,128 (71,292±13,139) | 866±49 (840±47) |
| | **M16** | 313 | 15 | 328 | 95.5 | 4.5 | 932 (988) | 0.30±0.05 (0.31±0.05) | 0.01±0.003 (0.01±0.003) | 0.31±0.05 (0.33±0.06) | 112,292±21,735 (104,431±20,214) | 88,399±17,309 (82,211±16,098) | 921±44 (894±43) |
| Vc | **AB2** | 475 | 23 | 498 | 95.4 | 4.6 | 1028 (1,056) | 0.41±0.06 (0.45±0.7) | 0.02±0.01 (0.02±0.01) | 0.44±0.06 (0.47±0.06) | 127,546±6,938 (118,618±6,453) | 82,215±4,379 (76,460±4,073) | 854±53 (828±51) |
| | **AB7** | 413 | 31 | 444 | 92.9 | 7.9 | 940 (963) | 0.39±0.05 (0.43±0.05) | 0.03±0.00 (0.03±0.00) | 0.42±0.05 (0.46±0.05) | 122,849±11,604 (114,249±10,792) | 85,620±9,319 (79,626±8,666) | 899±71 (872±69) |
| | **M16** | 369 | 33 | 402 | 92 | 8 | 1016 (1,099) | 0.33±0.11 (0.33±0.12) | 0.03±0.01 (0.03±0.01) | 0.35±0.12 (0.37±0.13) | 128,612±10,535 (119,474±9,797) | 94,297±9,401 (87,697±8,743) | 845±83 (819±80) |
| VI | **AB2** | 409 | 11 | 420 | 97.4 | 3.6 | 1020 (1,075) | 0.36±0.03 (0.38±0.03) | 0.01±0.00 (0.01±0.00) | 0.37±0.03 (0.39±0.03) | 110,144±8,402 (102,434±7,814) | 85,517±21,764 (79,531±20,241) | 861±71 (835±69) |
| | **AB7** | 361 | 17 | 378 | 95.2 | 4.8 | 1107 (1,130) | 0.29±0.04 (0.32±0.05) | 0.01±0.01 (0.02±0.01) | 0.31±0.04 (0.33±0.04) | 77,022±4,344 (71,630±4,040) | 38,043±10,184 (35,380±9,472) | 897±68 (870±66) |
| | **M16** | 407 | 25 | 432 | 94.2 | 5.8 | 992 (1,035) | 0.37±0.06 (0.40±0.07) | 0.02±0.01 (0.02±0.01) | 0.39±0.06 (0.42±0.07) | 111,303±5,379 (103,511±5,002) | 87,858±5,964 (81,708±5,547) | 796±79 (772±75) |

**Supplementary File 1d. Accumulated data acquired from the ultrastructural analysis of neuropil from layers I, II-is, II-ni, III, Va/b, Vc and VI of the MEC, for individual cases.** Data in parentheses are not corrected for shrinkage. AS: asymmetric synapses; CF: counting frame; SD: standard deviation; SE: standard error of the mean; SS: symmetric synapses.

| | AS | | | SS | | |
|---|---|---|---|---|---|---|
| | ***n*** | **M** | **σ** | ***n*** | **μ** | **σ** |
| **Layer I** | 1,253 | 11.34 | 0.71 | 80 | 10.65 | 0.71 |
| **Layer II-is** | 1,255 | 11.44 | 0.74 | 100 | 11.01 | 0.55 |
| **Layer II-ni** | 1,290 | 11.34 | 0.70 | 97 | 11.03 | 0.55 |
| **Layer III** | 1,210 | 11.48 | 0.78 | 74 | 10.97 | 0.75 |
| **Layer Va/b** | 1,143 | 11.60 | 0.69 | 50 | 11.04 | 0.61 |
| **Layer Vc** | 1,444 | 11.40 | 0.67 | 96 | 11.08 | 0.46 |
| **Layer VI** | 1,175 | 11.26 | 0.75 | 53 | 10.92 | 0.82 |
| **Layers I–VI** | 8,770 | 11.44 | 0.72 | 550 | 10.97 | 0.64 |

**Supplementary File 1e. Number of synaptic SAS analyzed (n), the location (μ) and scale (σ) of the best-fit log-normal distributions in the six cortical layers.** AS: asymmetric synapses; SAS: synaptic apposition surface; SS: symmetric synapses

| Layer | Type of Synapse | Macular | Perforated | Horseshoe-shaped | Fragmented | Complex |
|---|---|---|---|---|---|---|
| I | AS | 87% (1100) | 11.4% (143) | 1.3% (17) | 0.3% (4) | 13% (164) |
| | SS | 77.5% (62) | 15.1% (12) | 1.2% (1) | 6.2% (5) | 22.5% (18) |
| II-is | AS | 88.2% (1102) | 9% (113) | 2.2% (27) | 0.6% (8) | 11.8% (148) |
| | SS | 86.9% (86) | 8.1% (8) | 3% (3) | 2% (2) | 13.1% (13) |
| II-ni | AS | 86.3% (1110) | 11% (142) | 2.1% (27) | 0.6% (8) | 13.7% (177) |
| | SS | 73.2% (71) | 18.6% (18) | 7.2% (7) | 1% (1) | 26.8% (26) |
| III | AS | 78.9% (956) | 17.4% (211) | 2.7% (32) | 1% (12) | 21.1% (255) |
| | SS | 74.3% (55) | 18.9% (14) | 2.7% (2) | 4.1% (3) | 25.7% (19) |
| Va/b | AS | 81.6% (933) | 14.6% (167) | 2.6% (30) | 1.2% (14) | 18.4% (211) |
| | SS | 74% (37) | 14% (7) | 4% (2) | 8% (4) | 26% (13) |
| Vc | AS | 86.6% (1081) | 8.8% (110) | 3% (37) | 1.6% (20) | 13.4% (167) |
| | SS | 81.2% (69) | 11.8% (10) | 3.5% (3) | 3.5% (3) | 18.8% (16) |
| VI | AS | 77.6% (913) | 15.4% (181) | 3.8% (45) | 3.2% (37) | 22.4% (263) |
| | SS | 59.3% (32) | 25.9% (14) | 7.4% (4) | 7.4% (4) | 40.7% (22) |
| I-VI | AS | 83.9% (7195) | 12.4% (1067) | 2.5% (215) | 1.2% (103) | 16.1% (1.385) |
| | SS | 76.4% (412) | 15.4% (83) | 4.1% (22) | 4.1% (22) | 23.6% (127) |

**Supplementary File 1f. Proportion of the different shapes of synaptic junctions in MEC layers.** Data in parentheses refer to absolute numbers of synapses.

| Layer | Case | Type of Synapse | Macular | Perforated | Horseshoe-shaped | Fragmented | Complex |
|---|---|---|---|---|---|---|---|
| I | AB2 | AS | 88.4% (373) | 10.2% (43) | 0.9% (4) | 0.5% (2) | 11.6% (49) |
| | | SS | 74.2% (23) | 16.1% (5) | - (0) | 9.7% (3) | 25.8% (8) |
| | AB7 | AS | 89.6% (379) | 8.8% (37) | 1.6% (7) | - (0) | 10.4% (44) |
| | | SS | 78.2% (18) | 13% (3) | 4.4% (1) | 4.4% (1) | 21.8% (5) |
| | M16 | AS | 83.1% (348) | 15% (63) | 1.4% (6) | 0.5% (2) | 16.9% (71) |
| | | SS | 80.8% (21) | 15.4% (4) | - (0) | 3.8% (1) | 19.2% (5) |
| | Total | AS | 87% (1100) | 11.4% (143) | 1.3% (17) | 0.3% (4) | 13% (164) |
| | | SS | 77.5% (62) | 15.1% (12) | 1.2% (1) | 6.2% (5) | 22.5% (18) |
| II-is | AB2 | AS | 92.7% (392) | 6.2% (26) | 0.9% (4) | 0.2% (1) | 8.3% (31) |
| | | SS | 86.5% (32) | 8.1% (3) | 2.7% (1) | 2.7% (1) | 13.5% (5) |
| | AB7 | AS | 90.4% (386) | 6.1% (26) | 2.8% (12) | 0.7% (3) | 9.6% (41) |
| | | SS | 90% (27) | 6.7% (2) | - (0) | 3.3% (1) | 10% (3) |
| | M16 | AS | 81% (324) | 15.2% (61) | 2.8% (11) | 1% (4) | 19% (76) |
| | | SS | 84.4% (27) | 9.3% (3) | 6.3% (2) | - (0) | 15.6% (5) |
| | Total | AS | 88.2% (1102) | 9% (113) | 2.2% (27) | 0.6% (8) | 11.8% (148) |
| | | SS | 86.9% (86) | 8.1% (8) | 3% (3) | 2% (2) | 13.1% (13) |

| | | | | | | | |
|---|---|---|---|---|---|---|---|
| II-ni | AB2 | AS | 91.3% (390) | 7.3% (31) | 1.2% (5) | 0.2% (1) | 8.7% (37) |
| | | SS | 84.6% (22) | 3.8% (1) | 7.8% (2) | 3.8% (1) | 15.4% (4) |
| | AB7 | AS | 86.5% (418) | 10.4% (50) | 2.5% (12) | 0.6% (3) | 13.5% (65) |
| | | SS | 73.1% (19) | 23.1% (6) | 3.8% (1) | - (0) | 26.9% (7) |
| | M16 | AS | 80.1% (302) | 16.1% (61) | 2.7% (10) | 1.1% (4) | 19.9% (75) |
| | | SS | 66.7% (30) | 24.4% (11) | 8.9% (4) | - (0) | 33.3% (15) |
| | Total | AS | 86.3% (1110) | 11% (142) | 2.1% (27) | 0.6% (8) | 13.7% (177) |
| | | SS | 73.2% (71) | 18.6% (18) | 7.2% (7) | 1% (1) | 26.8% (26) |
| III | AB2 | AS | 80.8% (311) | 16.0% (62) | 1.6% (6) | 1.6% (6) | 19.2% (74) |
| | | SS | 71.4% (15) | 28.6% (6) | - (0) | - (0) | 28.6% (6) |
| | AB7 | AS | 80.7% (335) | 14.9% (62) | 3.7% (15) | 0.7% (3) | 19.3% (80) |
| | | SS | 85.7% (18) | 4.8% (1) | - (0) | 9.5% (2) | 14.3% (3) |
| | M16 | AS | 75.4% (310) | 21.2% (87) | 2.7% (11) | 0.7% (3) | 24.6% (101) |
| | | SS | 68.8% (22) | 21.9% (7) | 6.2% (2) | 3.1% (1) | 31.2% (10) |
| | Total | AS | 78.9% (956) | 17.4% (211) | 2.7% (32) | 1% (12) | 21.1% (255) |
| | | SS | 74.3% (55) | 18.9% (14) | 2.7% (2) | 4.1% (3) | 25.7% (19) |
| Va/b | AB2 | AS | 79.7% (333) | 18.1% (76) | 1% (4) | 1.2% (5) | 20.3% (85) |

| | | | | | | | |
|---|---|---|---|---|---|---|---|
| | | SS | 55.6% (10) | 33.2% (6) | 5.6% (1) | 5.6% (1) | 44.4% (8) |
| | AB7 | AS | 86.4% (357) | 9.4% (39) | 2.8% (12) | 1.2% (5) | 13.6% (56) |
| | | SS | 82.4% (14) | - (0) | - (0) | 17.6% (3) | 17.6% (3) |
| | M16 | AS | 77.6% (243) | 16.6% (52) | 4.5% (14) | 1.3% (4) | 22.4% (70) |
| | | SS | 86.6% (13) | 6.7% (1) | 6.7% (1) | - (0) | 13.4% (2) |
| | Total | AS | 81.6% (933) | 14.6% (167) | 2.6% (30) | 1.2% (14) | 18.4% (211) |
| | | SS | 74% (37) | 14% (7) | 4% (2) | 8% (4) | 26% (13) |
| Vc | AB2 | AS | 87.4% (415) | 8.6% (41) | 2.3% (11) | 1.7% (8) | 12.6% (60) |
| | | SS | 69.6% (16) | 26.1% (6) | 4.3% (1) | - (0) | 30.4% (7) |
| | AB7 | AS | 89.8% (368) | 6.6% (27) | 2.6% (11) | 1% (4) | 10.2% (42) |
| | | SS | 93.5% (29) | - (0) | 6.5% (2) | - (0) | 6.5% (2) |
| | M16 | AS | 82.1% (298) | 11.6% (42) | 4.1% (15) | 2.2% (8) | 17.9% (65) |
| | | SS | 77.4% (24) | 12.9% (4) | - (0) | 9.7% (3) | 22.6% (7) |
| | Total | AS | 86.6% (1081) | 8.8% (110) | 3% (37) | 1.6% (20) | 13.4% (167) |
| | | SS | 81.2% (69) | 11.8% (10) | 3.5% (3) | 3.5% (3) | 18.8% (16) |
| VI | AB2 | AS | 80.6% (329) | 11.5% (47) | 4.7% (19) | 3.2% (13) | 19.4% (79) |
| | | SS | 50% (6) | 33.3% (4) | 16.7% (2) | - (0) | 50% (6) |

| | | | | | | | |
|---|---|---|---|---|---|---|---|
| | AB7 | AS | 77.6% (280) | 18% (65) | 2.7% (10) | 1.7% (6) | 22.4% (81) |
| | | SS | 64.7% (11) | 23.5% (4) | 11.8% (2) | - (0) | 35.3% (6) |
| | M16 | AS | 74.7% (304) | 17% (69) | 3.9% (16) | 4.4% (18) | 25.3% (103) |
| | | SS | 60% (15) | 24% (6) | - (0) | 16% (4) | 40% (10) |
| | Total | AS | 77.6% (913) | 15.4% (181) | 3.8% (45) | 3.2% (37) | 22.4% (263) |
| | | SS | 59.3% (32) | 25.9% (14) | 7.4% (4) | 7.4% (4) | 40.7% (22) |
| I-VI | AB2 | AS | 86% (2543) | 11% (326) | 1.8% (53) | 1.2% (36) | 14% (415) |
| | | SS | 73.8% (124) | 18.5% (31) | 4.2% (7) | 3.6% (6) | 26.2% (44) |
| | AB7 | AS | 86.1% (2523) | 10.4% (306) | 2.7% (79) | 0.8% (24) | 13.9% (409) |
| | | SS | 82.4% (136) | 9.7% (16) | 3.7% (6) | 4.2% (7) | 17.6% (29) |
| | M16 | AS | 79.1% (2129) | 16.2% (435) | 3.1% (83) | 1.6% (43) | 20.9% (561) |
| | | SS | 73.7% (152) | 17.5% (36) | 4.4% (9) | 4.4% (9) | 26.3% (54) |
| | Total | AS | 83.9% (7195) | 12.4% (1067) | 2.5% (215) | 1.2% (103) | 16.1% (1385) |
| | | SS | 76.4% (412) | 15.4% (83) | 4.1% (22) | 4.1% (22) | 23.6% (127) |

**Supplementary File 1g. Proportion of the different shapes of synaptic junctions in MEC layers, for individual cases.** Data in parentheses refer to absolute numbers of synapses.

| Layer | Type of Synapse | Macular | Perforated | Horseshoe-Shaped | Fragmented | Complex |
|---|---|---|---|---|---|---|
| I | AS | 92,661 ± 3,095 (86,175 ± 2,878) | 197,654 ± 16,398 (183,819 ± 15,250) | 250,911 ± 40,834 (23,348 ± 37,975) | 267,410 ± 81,563 (248,691 ± 75,853) | 207,129 ± 20,454 (192,630 ±19,023) |
| | SS | 46,580 ± 4,067 (43,319 ± 3,782) | 79,066 ± 13,079 (73,531 ± 12,163) | 52,190 (48,537) | 66,841 ± 25,203 (62,163 ± 23,439) | 80,379 ± 14,474 (74,752 ± 13,461) |
| II-is | AS | 101,988 ± 7,669 (94,849 ± 7,133) | 247,884 ± 12,526 (230,532 ± 11,649) | 221,102 ± 35,576 (205,625 ± 33,086) | 206,647 ± 34,173 (192,182 ± 31,781) | 237,901 ± 12,971 (221,239 ± 12,065) |
| | SS | 65,361 ± 5,228 (60,785 ± 4,863) | 76,540 ± 22,208 (71,182 ± 20,653) | 61,474 ± 11,105 (57,171 ± 10,328) | 80,562 ± 17,663 (74,922 ± 16,426) | 72,531 ± 10,711 (67,454 ± 9,961) |
| II-ni | AS | 102,985 ± 4,837 (95,776 ± 4,499) | 208,141 ± 11,035 (193,571 ± 10,263) | 239,286 ± 20,749 (222,536 ± 19,297) | 210,844 ± 35,973 (196,085 ± 33,455) | 211,689 ± 10,498 (196,871 ± 9,763) |
| | SS | 63,792 ± 7,004 (59,327 ± 6,514) | 75,745 ± 17,776 (70,443 ± 16,531) | 90,677 ± 13,656 (84,330 ± 12,700) | 38,487 (35,793) | 81,962 ± 11,991 (76,224 ± 11,152) |
| III | AS | 98,632 ± 5,098 (91,728 ± 4,741) | 244,457 ± 10,273 (227,345 ± 9,554) | 221,461 ± 27,813 (205,958 ± 25,866) | 256,773 ± 29,314 (238,799 ± 27,262) | 244,603 ± 10,271 (227,481 ± 9,552) |
| | SS | 68,073 ± 9,960 (63,308 ± 9,263) | 89,039 ± 9,893 (82,806 ± 9,200) | 87,262 ± 43,491 (81,154 ± 40,447) | 150,035 ± 78,735 (139,532 ± 73,223) | 90,822 ± 11,898 (84,465 ±11,065) |

| | | | | | | |
|---|---|---|---|---|---|---|
| **Va/b** | **AS** | 112,898 ± 4,214 (104,995 ± 3,919) | 242,129 ± 11,233 (225,180 ± 10,447) | 220,571 ± 11,053 (205,131 ± 10,279) | 253,021 ± 32,899 (235,310 ± 30,596) | 238,420 ± 9,232 (221,731 ± 8,586) |
| | **SS** | 65,199 ± 10,194 (60,635 ± 9,481) | 82,702 ± 8,009 (76,913 ± 7,448) | 40,590 ± 886 (37,749± 824) | 122,609 ± 15,686 (114,026 ± 14,588) | 91,303 ± 14,198 (84,911 ± 13,204) |
| **Vc** | **AS** | 108,431 ± 5,220 (100,841 ± 4,854) | 240,291 ± 17,494 (223,470 ± 16,269) | 243,754 ± 25,012 (226,691 ± 23,262) | 265,690 ± 24,100 (247,092 ± 22,413) | 108,431 ± 5,220 (100,841 ± 4,854) |
| | **SS** | 83,176 ± 5,150 (77,354 ± 4,790) | 111,784 ± 13,144 (103,959 ± 12,224) | 73,753 ± 17,089 (68,590 ± 15,892) | 101,698 ± 15,405 (94,580 ± 14,327) | 83,176 ± 5,150 (77,354 ± 4,790) |
| **VI** | **AS** | 79,331 ± 4,767 (73,778 ± 4,433) | 168,558 ± 15,001 (156,759 ± 13,951) | 183,334 ± 14,939 (170,501 ± 13,893) | 191,595 ± 19,309 (178,183 ± 17,957) | 170,959 ± 12,388 (158,992 ± 11,521) |
| | **SS** | 44,170 ± 6,484 (41,078 ± 6,030) | 119,142 ± 13,275 (110,802 ± 12,346) | 62,251 ± 17,584 (57,894 ± 16,354) | 129,831 ± 35,440 (120,742 ± 32,959) | 110,547 ± 13,584 (102,808 ± 12,634) |
| **I-VI** | **AS** | 99,561 ± 2,260 (92,592 ± 2,101) | 221,302 ± 6,078 (205,811 ± 5,653) | 225,369 ± 9,858 (209,593 ± 9,168) | 233,668 ± 11,949 (217,311 ± 11,112) | 99,561 ± 2,260 (92,592 ± 2,101) |
| | **SS** | 62,336 ± 3,029 (57,972 ± 2,817) | 91,444 ± 5,739 (85,043 ± 5,337) | 70,680 ± 7,324 (65,733 ± 6,811) | 102,349 ± 15,452 (95,185 ± 14,371) | 62,336 ± 3,029 (57,972 ± 2,817) |

**Supplementary File 1h. SAS area of asymmetric (AS) and symmetric (SS) synapses for each synaptic shape, in each MEC layer.** Data in parentheses are not corrected for shrinkage. Values (in $nm^2$) are expressed as mean ± standard error.

| Layer | Case | Tipe of Synapse | Macular | Perforated | Horseshoe | Fragmented | Complex |
|---|---|---|---|---|---|---|---|
| I | AB2 | AS | 101,222 ± 3,039 (94,136 ± 2,827) | 184,880 ± 29,944 (171,938 ± 27,848) | 175,065 ± 59,004 (162,810 ± 54,874) | 322,942 ± 103,471 (300,036 ± 96,228) | 191,851 ± 34,956 (178,422 ± 32,509) |
| | | SS | 38,842 ± 2,120 (36,123 ± 1,972) | 59,168 ± 5,204 (55,026 ± 4,840) | - | 85,613 (79,620) | 90,854 ± 36,558 (84,494 ± 33,999) |
| | AB7 | AS | 89,966 ± 3,225 (83,669 ± 2,999) | 212,981 ± 44,328 (198,072 ± 41,225) | 348,624 ± 72,012 (324,220 ± 66,971) | - | 235,190 ± 54,498 (218,727 ± 50,683) |
| | | SS | 55,430 ± 1,433 (51,550 ± 1,333) | 77,980 ± 27,699 (72,521 ± 25,760) | 52,190 (48,537) | 36,352 (33,808) | 66,743 ± 19,545 (62,071 ± 18,177) |
| | M16 | AS | 86,795 ± 6,152 (80,720 ± 5,721) | 195,103 ± 12,753 (181,446 ± 11,860) | 203,763 ± 37,594 (189,500 ± 34,962) | 156,347 (145,403) | 194,345 ± 15,380 (180,741 ± 14,303) |
| | | SS | 45,467 ± 11,054 (42,285 ± 10,280) | 100,051 ± 30,562 (93,048 ± 28,423) | - | 41,015 (38,144) | 85,118 ± 15,629 (79,160 ± 14,535) |
| II-is | AB2 | AS | 130,221 ± 5,857 (121,105 ± 5,447) | 286,237 ± 21,844 (266,201 ± 20,315) | 300,127 ± 94,855 (279,118 ± 88,215) | 175,889 (163,577) | 278,763 ± 26,510 (259,250 ± 24,655) |
| | | SS | 73,872 ± 7,156 (68,701 ± 6,655) | 141,358 (131,463) | 50,369 (46,844) | 62,899 (58,496) | 86,056 ± 35,687 (80,032 ± 33,189) |
| | AB7 | AS | 89,423 ± 4,109 (83,164 ± 3,821) | 232,829 ± 15,903 (216,531 ± 14,790) | 183,706 ± 37,076 (170,847 ± 34,480) | 166,100 ± 36,220 (154,473 ± 33,685) | 216,045 ± 6,865 (200,922 ± 6,385) |
| | | SS | 53,519 ± 3,588 (49,773 ± 3,337) | 40,445 (37,614) | - | 98,224 (91,348) | 69,335 ± 28,890 (64,481 ± 26,867) |
| | M16 | AS | 86,321 ± 7,379 (80,279 ± 6,863) | 224,586 ± 5,709 (208,865 ± 5,309) | 179,473 ± 11,288 (166,910 ± 10,498) | 282,845 ± 69,543 (263,046 ± 64,675) | 218,864 ± 3,840 (203,544 ± 3,571) |
| | | SS | 68,691 ± 12,332 (63,882 ± 11,469) | 62,178 ± 868 (58,095 ± 808) | 72,579 (67,499) | - | 65,645 ± 3,503 (61,050 ± 3,258) |

| | | | | | | | |
|---|---|---|---|---|---|---|---|
| II-ni | AB2 | AS | 114,835 ± 6,400 (106,796 ± 5,952) | 211,495 ± 24,500 (196,690 ± 22,785) | 269,605 ± 15,871 (250,733 ± 56,240) | 273,833 (254,665) | 222,354 ± 29,133 (205,118 ± 27,093) |
| | | SS | 75,232 ± 13,131 (69,966 ± 12,211) | 45,204 (42,040) | 117,137 (108,937) | 38,487 (35,793) | 65,823 ± 27,336 (61,215 ± 25,423) |
| | AB7 | AS | 90,025 ± 8,873 (83,723 ± 8,252) | 189,558 ± 17,745 176,289 (16,503) | 224,039 ± 9,971 (208,356 ± 9,273) | 281,632 ± 46,258 (261,918 ± 43,020) | 196,410 ± 10,908 (179,157 ± 10,144) |
| | | SS | 41,902 ± 4,454 (38,969 ± 4,143) | 74,983 ± 30,956 (69,734 ± 28,790) | 71,960 (66,923) | - | 79,189 ± 28,356 (73,646 ± 26,371) |
| | M16 | AS | 104,096 ± 2,504 (96,809 ± 2,329) | 223,371 ± 15,871 (207,735 ± 16,503) | 224,214 ± 26,844 (208,519 ± 24,965) | 142,656 ± 33,174 (132,670 ± 30,852) | 216,305 ± 13,061 (208,541 ± 12,146) |
| | | SS | 74,241 ± 6072 (69,044 ± 5,647) | 86,688 ± 32,217 (80,620 ± 29,962) | 86,806 ± 24,058 (80,730 ± 22,374) | - | 95,494 ± 10,325 (88,809 ± 9,602) |
| III | AB2 | AS | 110,929 ± 7,912 (103,164 ± 7,358) | 247,376 ± 8,401 (230,059 ± 7,813) | 257,345 ± 62,238 (239,331 ± 57,881) | 295,531 ± 37,375 (274,844 ± 34,759) | 253,575 ± 4,765 (235,825 ± 4,431) |
| | | SS | 101,194 ± 14,339 (94,111 ± 13,335) | 82,440 ± 3,302 (76,669 ± 2,820) | - | - | 82,440 ± 3,032 (76,669 ± 2,820) |
| | AB7 | AS | 93,165 ± 6,029 (86,644 ± 5,607) | 250,210 ± 32,209 (232,695 ± 9,954) | 246,375 ± 30,086 (229,129 ± 27,980) | 157,049 ± 26,305 (146,055 ± 24,463) | 247,938 ± 32,422 (230,582 ± 30,152) |
| | | SS | 49,072 ± 10,065 (45,637 ± 9,361) | 45,371 (42,195) | - | 71,687 ± 13,513 (66,669 ± 12,568) | 61,730 ± 3,556 (57,409 ± 3,307) |
| | M16 | AS | 91,802 ± 9,950 (85,376 ± 9,254) | 235,785 ± 10,000 (219,280 ± 9,300) | 160,662 ± 41,252 (149,415 ± 38,364) | 284,496 ± 44,073 (264,582 ± 40,988) | 232,297 ± 8,405 (216,037 ± 7,817) |

| | | | | | | | |
|---|---|---|---|---|---|---|---|
| | | SS | 53,953 ± 7,397 (50,176 ± 6,880) | 110,193 ± 10,607 (102,480 ± 9,865) | 87,262 ± 43,491 (81,154 ± 40,447) | 306,729 (285,258) | 118,599 ± 24,592 (110,297 ± 22,871) |
| Va/b | AB2 | AS | 123,697 ± 5,048 (115,038 ± 4,695) | 236,015 ± 13,588 (219,494 ± 12,637 | 241,207 ± 28,095 (224,323 ± 26,128) | 325,348 ± 8,295 (302,574 ± 7,714) | 242,546 ± 9,149 (225,568 ± 8,509) |
| | | SS | 38,564 ± 13,183 (35,865 ± 12,260) | 77,504 ± 8,616 (72,078 ± 8,013) | 39,704 (36,925) | 92,963 (86,456) | 74,827 ± 10,849 (69,589 ± 10,090) |
| | AB7 | AS | 100,378 ± 4,258 (93,351 ± 3,960) | 234,281 ± 14,117 (217,882 ± 12,637) | 207,968 ± 15,552 (193,410 ± 14,463) | 238,907 ± 69,200 (222,183 ± 64,356) | 229,614 ± 14,206 (213,541 ± 13,211) |
| | | SS | 67,421 ± 9,282 (62,702 ± 8,632) | - | - | 137,432 ± 8,890 (127,812 ± 8,268) | 137,432 ± 8,890 (127,812 ± 8,268) |
| | M16 | AS | 114,620 ± 5,597 (106,596 ± 5,205) | 256,091 ± 31,359 (238,165 ± 29,164) | 212,538 ± 10,497 (197,661 ± 9,762) | 158,645 ± 27,959 (147,540 ± 26,002) | 243,100 ± 26,059 (226,083 ± 24,235) |
| | | SS | 89,611 ± 18,220 (83,338 ± 16,945) | 98,297 (91,416) | 41,476 (38,573) | - | 69,887 ± 28,410 (64,995 ± 26,421) |
| Vc | AB2 | AS | 108,768 ± 2,541 (101,155 ± 2,363) | 245,585 ± 22,685 (228,394 ± 21,097) | 302,117 ± 48,067 (280,969 ± 44,703) | 250,032 ± 32,883 (232,530 ± 30,581) | 250,483 ± 23,052 (225,568 ± 8,509) |
| | | SS | 71,875 ± 811 (66,844 ± 754) | 117,148 ± 6,131 (108,948 ± 5,702) | 74,859 (69,619) | - | 111,096 ± 78 (103,310 ± 73) |
| | AB7 | AS | 111,861 ± 11,060 (104,031 ± 10,286) | 221,124 ± 5,818 (205,645 ± 5,411) | 221,937 ± 41,817 (206,401 ± 38,890) | 231,859 ± 79,721 (215,629 ± 74,141) | 223,918 ± 19,853 (213,541 ± 13,211) |
| | | SS | 86,082 ± 8,400 (80,056 ± 7,812) | - | 73,200 ± 29,583 (68,076 ± 27,512) | - | 73,200 ± 29,583 (8,076 ± 27,512) |

| | | | | | | | |
|---|---|---|---|---|---|---|---|
| | M16 | AS | 104,665 ± 13,607 (97,338 ± 12,655) | 254,163 ± 53,198 (236,371 ± 49,474) | 207,207 ± 28,972 (192,703 ± 26,944) | 303,903 ± 31,417 (282,630 ± 29,218) | 244,450 ± 39,981 (227,338 ± 37,182) |
| | | SS | 91,570 ± 11,990 (85,160 ± 11,151) | 108,207 ± 23,396 (100,633 ± 21,758) | - | 101,698 ± 15,405 (94,580 ± 14,327) | 108,844 ± 32,951 (101,225 ± 16,404) |
| VI | AB2 | AS | 86,228 ± 6,132 (80,192 ± 5,703) | 220,662 ± 15,119 (205,216 ± 14,060) | 177,145 ± 38,427 (164,745 ± 35,737) | 224,731 ± 42,881 (209,000 ± 39,879) | 209,198 ± 8,633 (194,554 ± 8,029) |
| | | SS | 39,253 ± 5,372 (36,506 ± 4,996) | 130,164 ± 40,331 (121,053 ± 37,508) | 94,885 (88,243) | - | 131,427 ± 39,068 (122,227 ± 36,333) |
| | AB7 | AS | 63,184 ± 3,185 (58,761 ± 2,962) | 121,789 ± 2,559 (113,264 ± 2,380) | 165,155 ± 26,120 (153,594 ± 24,291) | 144,688 ± 27,602 (134,560 ± 25,669) | 126,350 ± 3,346 (117,506 ± 3,111) |
| | | SS | 28,110 ± 5,038 (26,143 ± 4,686) | 90,385 ± 9,873 (84,058 ± 9,182) | 45,935 ± 11,354 (42,719 ± 10,560) | - | 71,063 ± 3,643 (66,088 ± 3,388) |
| | M16 | AS | 88,482 ± 5,290 (82, 382 ± 4,920) | 163,224 ± 185 (151,798 ± 173) | 207,702 ± 6,165 (193,163 ± 5,734) | 205,366 ± 11,372 (190,990 ± 10,576) | 177,328 ± 3,086 (164,915 ± 2,870) |
| | | SS | 65,146 ± 9,449 (60,586 ± 8,787) | 130,965 ± 16,519 (121,798 ± 5,362) | - | 129,831 ± 35,440 (120,742 ± 32,959) | 122,949 ± 6,055 (114,342 ± 5,631) |

**Supplementary File 1i. SAS area of asymmetric (AS) and symmetric (SS) synapses for each synaptic shape, in each MEC layer, per individual case.** Data in parentheses are not corrected for shrinkage. Values (in $nm^2$) are expressed as mean ± standard error.

| Layer | Type of Synapse | Spine Head | Spine Neck | Spiny Shaft | Aspiny Shaft | Shafts (Spiny + Aspiny) |
|---|---|---|---|---|---|---|
| I | AS | 54.2% (675) | 0.4% (5) | 17.5% (218) | 21.9% (273) | 39.4% (491) |
| | SS | 0.6% (7) | 0.2% (3) | 1.8% (23) | 3.4% (43) | 5.2% (66) |
| II-is | AS | 57.5% (740) | 0.6% (8) | 15.7% (202) | 18.5% (240) | 34.2% (442) |
| | SS | 1.1% (14) | 0.2% (2) | 3.9% (50) | 2.5% (32) | 6.4% (82) |
| II-ni | AS | 59.5% (798) | 0.2% (3) | 16.2% (217) | 17.1% (229) | 33.3% (446) |
| | SS | 0.5% (7) | 0.1% (2) | 4.2% (56) | 2.2% (30) | 6.4% (86) |
| III | AS | 54.9% (667) | 0.5% (6) | 20.9% (255) | 17.9% (218) | 38.8% (473) |
| | SS | 0.7% (9) | - (0) | 3.0% (37) | 2.1% (26) | 5.1% (63) |
| Va/b | AS | 52.9% (571) | 1.0% (11) | 21.5% (232) | 20.9% (226) | 42.4% (458) |
| | SS | 0.1% (1) | 0.2% (2) | 1.8% (19) | 1.6% (17) | 3.4% (36) |
| Vc | AS | 60.5% (761) | 0.7% (9) | 14.5% (183) | 17.8% (223) | 32.3% (406) |
| | SS | 1.4% (18) | 0.1% (1) | 2.6% (33) | 2.4% (30) | 5% (63) |
| VI | AS | 76.2% (850) | 0.3% (3) | 9.4% (106) | 9.8% (109) | 19.2% (215) |
| | SS | 0.6% (6) | 0.1% (1) | 2.2% (24) | 1.4% (16) | 3.6% (40) |
| I-VI | AS | 59.3% (5062) | 0.5% (45) | 16.5% (1413) | 17.8% (1518) | 34.3% (2931) |
| | SS | 0.7% (62) | 0.1% (11) | 2.8% (242) | 2.3% (194) | 5.1% (436) |

**Supplementary File 1j. Proportion of the different postsynaptic targets in MEC layers.** Synapses established on spine head include both complete and incomplete spines. Data in parentheses refer to the absolute number of synapses found in each layer.

| Layer | Case | Type of Synapse | Complete Spine Heads | Incomplete Spine Heads | Spine Necks | Spiny Shaft | Aspiny Shaft | Shafts (Spiny + Aspiny) |
|---|---|---|---|---|---|---|---|---|
| I | AB2 | AS | 41.6% (160) | 13.0% (50) | 0.3% (1) | 18% (69) | 27.1% (104) | 45.1% (173) |
| | | SS | - (0) | - (0) | 3.5% (1) | 17.2% (5) | 79.3% (23) | 96.5% (27) |
| | AB7 | AS | 39.6% (156) | 17.6% (69) | 1% (4) | 17.6% (63) | 24.2% (95) | 41.8% (158) |
| | | SS | - (0) | 23.8% (5) | 4.8% (1) | 33.3% (7) | 38.1% (8) | 71.4% (15) |
| | M16 | AS | 40.1% (158) | 20.8% (82) | - (0) | 20.3% (80) | 18.8% (74) | 39.1% (154) |
| | | SS | 3.8% (1) | 3.8% (1) | 3.8% (1) | 42.4% (11) | 46.2% (12) | 88.6% (23) |
| | Total | AS | 40.5% (474) | 17.2% (201) | 0.4% (5) | 18.6% (218) | 23.3% (273) | 41.9% (491) |
| | | SS | 1.3% (1) | 7.9% (6) | 3.9% (3) | 30.3% (23) | 56.6% (43) | 86.9% (66) |
| II-is | AB2 | AS | 38.5% (155) | 17.4% (70) | 0.2% (1) | 14.4% (58) | 29.5% (119) | 43.9% (177) |
| | | SS | 10.8% (4) | - (0) | 2.7% (1) | 51.4% (19) | 35.1% (13) | 86.5% (32) |
| | AB7 | AS | 49% (200) | 12.3% (69) | 1.2% (4) | 20.1% (82) | 17.4% (71) | 37.5% (153) |
| | | SS | 13.8% (4) | - (0) | - (0) | 48.3% (14) | 37.9% (11) | 86.2% (35) |
| | M16 | AS | 48.8% (185) | 21.1% (80) | 0.5% (2) | 16.4% (60) | 13.2% (50) | 29.6% (110) |
| | | SS | 15.6% (5) | 3.1% (1) | 3.1% (1) | 53.2% (17) | 25% (8) | 78.2% (25) |
| | Total | AS | 45.3% (540) | 16.8% (201) | 0.7% (7) | 17.0% (202) | 20.2% (240) | 37.2% (442) |
| | | SS | 13.3% (13) | 1.0% (1) | 2.0% (2) | 51.0% (50) | 32.7% (32) | 83.7% (82) |

| | | | | | | | | |
|---|---|---|---|---|---|---|---|---|
| **II-ni** | **AB2** | **AS** | 36.9% (155) | 20.2% (85) | 0.2% (1) | 18.7% (78) | 24% (101) | 42.7% (179) |
| | | **SS** | 3.8% (1) | - (0) | - (0) | 42.4% (11) | 53.8% (14) | 96.2% (35) |
| | **AB7** | **AS** | 47.8% (220) | 17.2% (79) | 0.2% (1) | 18.7% (86) | 16.1% (74) | 34.8% (160) |
| | | **SS** | 15.4% (4) | - (0) | 3.8% (1) | 38.5% (10) | 42.3% (11) | 80.8% (21) |
| | **M16** | **AS** | 48.8% (202) | 21.1% (57) | 0.5% (1) | 16.4% (54) | 13.2% (53) | 29.6% (107) |
| | | **SS** | 2.3% (1) | 2.3% (1) | 2.3% (1) | 81.5% (35) | 11.6% (5) | 93.1% (40) |
| | **Total** | **AS** | 46.3% (577) | 17.7% (221) | 0.2% (3) | 17.4% (217) | 18.4% (229) | 35.8% (446) |
| | | **SS** | 6.3% (6) | 1.1% (1) | 2.1% (2) | 58.9% (56) | 31.6% (30) | 90.5% (86) |
| **III** | **AB2** | **AS** | 37.5% (135) | 21.4% (77) | - (0) | 21.4% (77) | 19.7% (71) | 41.1% (148) |
| | | **SS** | 5% (1) | - (0) | - (0) | 55% (11) | 40% (8) | 95% (19) |
| | **AB7** | **AS** | 39.5% (153) | 18.1% (70) | 1% (3) | 19.2% (74) | 22.2% (86) | 41.4% (160) |
| | | **SS** | 10% (2) | 10% (2) | - (0) | 20% (4) | 60% (12) | 80% (16) |
| | **M16** | **AS** | 45.5% (179) | 13.5% (53) | 0.8% (3) | 27.2% (107) | 13% (51) | 40.2% (158) |
| | | **SS** | 9.4% (3) | 3.1% (1) | - (0) | 68.7% (22) | 18.8% (6) | 87.5% (28) |
| | **Total** | **AS** | 40.8% (467) | 17.5% (200) | 0.5% (6) | 22.2% (255) | 19% (218) | 41.2% (473) |
| | | **SS** | 8.3% (6) | 4.2% (3) | - (0) | 51.4% (37) | 36.1% (26) | 87.5% (63) |
| **Va/b** | **AB2** | **AS** | 37.5% (140) | 16.9% (63) | 1.9% (7) | 23.9% (89) | 19.8% (74) | 43.7% (163) |

| | | | | | | | | |
|---|---|---|---|---|---|---|---|---|
| | | **SS** | 7.1% (1) | - (0) | 7.1% (1) | 64.4% (9) | 21.4% (3) | 85.8% (12) |
| | **AB7** | **AS** | 46% (180) | 15.4% (60) | 0.5% (2) | 21.7% (85) | 16.4% (64) | 38.1% (149) |
| | | **SS** | - (0) | - (0) | 6.3% (1) | 43.7% (4) | 50.0% (12) | 93.7% (16) |
| | **M16** | **AS** | 24.3% (67) | 22.1% (61) | 0.7% (2) | 21% (58) | 31.9% (88) | 52.9% (146) |
| | | **SS** | - (0) | - (0) | - (0) | 33.3% (3) | 66.7% (6) | 100% (9) |
| | **Total** | **AS** | 37.2% (387) | 17.7% (184) | 1.1% (11) | 22.3% (232) | 21.7% (226) | 44% (458) |
| | | **SS** | 2.6% (1) | - (0) | 5.1% (2) | 48.7% (19) | 43.6% (17) | 92.3% (36) |
| **Vc** | **AB2** | **AS** | 31.5% (142) | 33.8% (153) | 0.7% (3) | 12.9% (58) | 21.1% (95) | 34% (153) |
| | | **SS** | 9.1% (2) | 9.1% (2) | - (0) | 36.4% (8) | 45.4% (10) | 81.8% (18) |
| | **AB7** | **AS** | 40.8% (154) | 27.6% (104) | 0.8% (3) | 20.7% (78) | 10.1% (38) | 30.8% (116) |
| | | **SS** | 16.7% (5) | 3.3% (1) | - (0) | 46.7% (14) | 33.3% (10) | 80% (24) |
| | **M16** | **AS** | 35.6% (124) | 24.1% (84) | 0.9% (3) | 13.5% (47) | 25.9% (90) | 39.4% (137) |
| | | **SS** | 26.7% (8) | - (0) | 3.3 (1) | 36.7% (11) | 33.3% (10) | 70% (21) |
| | **Total** | **AS** | 35.7% (420) | 29% (341) | 0.8% (9) | 15.5% (183) | 19% (223) | 34.5% (406) |
| | | **SS** | 18.2% (15) | 3.6% (3) | 1.2% (1) | 40.3% (33) | 36.7% (30) | 77% (63) |
| **Vl** | **AB2** | **AS** | 46.9% (167) | 31.2% (111) | 0.6% (2) | 11.8% (42) | 9.5% (34) | 21.3% (76) |
| | | **SS** | - (0) | - (0) | - (0) | 45.5% (5) | 54.5% (10) | 100% (15) |

| | | | | | | | | |
|---|---|---|---|---|---|---|---|---|
| | **AB7** | **AS** | 47.5% (164) | 29.0% (100) | - (0) | 9.3% (32) | 14.2% (49) | 23.5% (81) |
| | | **SS** | 13.3% (2) | 6.7% (1) | - (0) | 66.7% (10) | 13.3% (2) | 80% (12) |
| | **M16** | **AS** | 50.1% (184) | 33.8% (124) | 0.3% (1) | 8.7% (32) | 7.1% (26) | 15.8% (58) |
| | | **SS** | - (0) | 14.2% (3) | 4.8% (1) | 42.9% (9) | 38.1% (8) | 81% (17) |
| | **Total** | **AS** | 48.2% (515) | 31.4% (335) | 0.3% (3) | 9.9% (106) | 10.2% (109) | 20.1% (215) |
| | | **SS** | 4.3% (2) | 8.5% (4) | 2.1% (1) | 51.1% (24) | 34% (16) | 85.1% (40) |
| **I-VI** | **AB2** | **AS** | 38.4% (1054) | 22.2% (609) | 0.5% (15) | 17.1% (471) | 21.8% (598) | 38.9% (1069) |
| | | **SS** | 5.7% (9) | 1.3% (2) | 1.8% (3) | 42.8% (68) | 48.4% (77) | 91.2% (145) |
| | **AB7** | **AS** | 44.3% (1227) | 19.2% (532) | 0.7% (18) | 18.2% (503) | 17.6% (487) | 35.8% (990) |
| | | **SS** | 10.8% (17) | 5.7% (9) | 1.9% (3) | 42% (66) | 39.6% (62) | 81.6% (128) |
| | **M16** | **AS** | 43.5% (1099) | 21.4% (541) | 0.5% (12) | 17.4% (439) | 17.2% (433) | 34.6% (872) |
| | | **SS** | 9.3% (18) | 3.6% (7) | 2.6% (5) | 56% (108) | 28.5% (55) | 84.5% (163) |
| | **Total** | **AS** | 42.1% (3380) | 20.9% (1682) | 0.5% (45) | 17.6% (1413) | 18.9% (1518) | 36.5% (2931) |
| | | **SS** | 8.6% (44) | 3.5% (18) | 2.2% (11) | 47.5% (242) | 38.2% (194) | 85.7% (436) |

**Supplementary File 1k. Proportion of the post-synaptic targets of synaptic junctions in MEC layers, for individual cases**. Total synapses on spine heads are calculated as the sum of “Complete Spine Heads” and “Incomplete Spine Heads” columns. Data in parentheses refer to the absolute numbers of synapses.

| Layer | Type of Synapse | Spine Head | Spine Neck | Spiny Shaft | Aspiny Shaft | Shafts (Spiny + Aspiny) |
|---|---|---|---|---|---|---|
| I | AS | 119,082 ± 6,100 (110,747 ± 5,673) | 68,168 ± 18,563 (62,396 ± 17,263) | 101,986 ± 5,371 (94,847 ± 4,995) | 98,581 ± 4,375 (91,680 ± 4,069) | 99,960 ± 3,545 (92,962 ± 3,297) |
| | SS | 161,971 (150,633) | 42,755 ± 3,477 (39,762 ± 3,234) | 48,287 ± 6,885 (44,907 ± 6,403) | 46,804 ± 3,553 (43,528 ± 3,305) | 48,094 ± 2,042 (44,727 ± 1,899) |
| II-is | AS | 125,834 ± 6,599 (117,025 ± 6,137) | 78,709 ± 20,174 (73,200 ± 18,761) | 117,234 ± 6,063 (109,028 ± 5,639) | 110,359 ± 7,633 (102,634 ± 7,098) | 115,956 ± 5,572 (107,839 ± 5,182) |
| | SS | 68,081 ± 11,271 (63,316 ± 10,482) | 70,880 ± 4,112 (65,919 ± 3,824) | 72,899 ± 7,931 (67,796 ± 7,376) | 67,660 ± 6,457 (62,924 ± 6,005) | 69,017 ± 5,549 (64,186 ± 5,160) |
| II-ni | AS | 124,614 ± 8,464 (115,891 ± 7,872) | 69,252 ± 3,827 (64,404 ± 3,559) | 138,955 ± 6,692 (129,228 ± 6,224) | 108,619 ± 8,056 (101,016 ± 7,493) | 124,575 ± 6,706 (115,854 ± 6,237) |
| | SS | 58,746 ± 25,087 (54,633 ± 23,331) | 43,048 ± 5,728 (40,035 ± 5,327) | 74,894 ± 9,012 (69,651 ± 8,381) | 72,447 ± 10,324 (67,376 ± 9,601) | 77,563 ± 8,752 (72,134 ± 8,140) |
| III | AS | 143,154 ± 7,462 (133,133 ± 6,940) | 69,862 ± 18,204 (64,972 ± 16,929) | 139,844 ± 8,879 (130,055 ± 8,258) | 127,158 ± 8,187 (118,257 ± 7,641) | 131,819 ± 7,258 (122,591 ± 6,750) |
| | SS | 112,999 ± 46,011 (105,089 ± 42,791) | - | 82,262 ± 11,026 (76,503 ± 10,255) | 66,073 ± 9,339 (61,448 ± 8,685) | 72,577 ± 8,570 (67,496 ± 7,970) |
| Va/b | AS | 148,730 ± 7,348 (138,319 ± 7,348) | 67,245 ± 16,663 (62,537 ± 15,497) | 151,189 ± 10,398 (140,606 ± 9,670) | 137,209 ± 5,457 (127,604 ± 5,075) | 143,122 ± 6,939 (133,104 ± 6,453) |
| | SS | 71,678 (66,660) | 32,816 ± 20,518 (30,519 ± 19,082) | 59,604 ± 5,089 (55,432 ± 4,733) | 83,131 ± 12,141 (77,312 ± 11,291) | 67,851 ± 6,157 (63,101 ± 5,726) |
| Vc | AS | 128,645 ± 6,101 (119,346 ± 5,673) | 60,443 ± 8,584 (56,212 ± 7,983) | 135,639 ± 5,381 (126,144 ± 5,004) | 131,950 ± 7,869 (122,714 ± 7,319) | 127,584 ± 5,012 (118,653 ± 4,661) |

| | | | | | | |
|---|---|---|---|---|---|---|
| | **SS** | 79,588 ± 10,548 (74,017 ± 9,809) | 115,738 (107,636) | 96,546 ± 10,923 (89,788 ± 10,158) | 86,027 ± 4,670 (80,005 ± 4,343) | 85,941 ± 6,115 (79,926 ± 5,687) |
| **VI** | **AS** | 105,948 ± 8,586 (98,532 ± 7,985) | 48,950 ± 6,519 (45,524 ± 6,063) | 99,751 ± 12,528 (92,769 ± 11,651) | 105,833 ± 10,656 (98,424 ± 9,910) | 103,833 ± 9,881 (96,564 ± 9,189) |
| | **SS** | 33,254 ± 18,289 (30,918 ± 17,009) | 52,193 (48,540) | 96,837 ± 11,956 (90,058 ± 11,119) | 51,591 ± 11,093 (47,980 ± 10,317) | 70,343 ± 10,522 (65,419 ± 9,786) |
| **I-VI** | **AS** | 127,956 ± 3,120 (118,999 ± 2,902) | 66,435 ± 5,616 (61,784 ± 5,223) | 126,371 ± 3,808 (117,525 ± 3,542) | 117,101 ± 3,266 (108,904 ± 3,038) | 120,978 ± 3,006 (112,510 ± 2,796) |
| | **SS** | 76,801 ± 9,326 (71,425 ± 8,673) | 53,608 ± 7,928 (49,855 ± 7,373) | 76,403 ± 3,959 (71,055 ± 3,681) | 68,026 ± 3,560 (63,264 ± 3,311) | 70,198 ± 2,948 (65,284 ± 2,742) |

**Supplementary File 1l. SAS area of asymmetric (AS) and symmetric (SS) synapses regarding the post-synaptic target, in each MEC layer**. Data in parentheses are not corrected for shrinkage. Values (in $nm^2$) are expressed as mean ± standard error.

| Layer | Case | Type of Synapse | Spine Head | Spine Neck | Spiny Shaft | Aspiny Shaft | Shafts (Spiny + Aspiny) |
|---|---|---|---|---|---|---|---|
| I | AB2 | AS | 120,467 ± 9,430 (112,034 ± 8,770) | 92,839 (86,340) | 118,246 ± 6,221 (109,969 ± 5,785) | 97,256 ± 8,078 (90,448 ± 3,157) | 105,655 ± 2,639 (98,259 ± 2,454) |
| | | SS | - | 43,604 (40,552) | 43,049 ± 16,837 (40,035 ± 15,658) | 45,416 ± 1,848 (42,237 ± 1,719) | 47,413 ± 2,243 (44,094 ± 2,086) |
| | AB7 | AS | 115,304 ± 7,209 (107,232 ± 6,705) | 55,833 ± 24,026 (51,924 ± 2,344) | 89,724 ± 5,185 (83,444 ± 4,822) | 100,994 ± 9,662 (93,924 ± 8,986) | 96,381 ± 5,605 (89,634 ± 5,213) |
| | | SS | - | 36,352 (33,808) | 52,413 ± 8,390 (48,744 ± 7,803) | 50,674 ± 6,169 (47,127 ± 5,737) | 49,327 ± 5,114 (45,874 ± 4,756) |
| | M16 | AS | 121,477 ± 17,164 (108,683 ± 15,962) | - | 97,989 ± 8,078 (91,130 ± 7,513) | 97,493 ± 10,973 (90,669 ± 10,205) | 97, 843 ± 9,358 (90,994 ± 8,703) |
| | | SS | 161,971 (150,633) | 48,307 (44,926) | 52,017 ± 1,577 (48,376 ± 1,467) | 44,322 ± 9,926 (41,219 ± 9,231) | 47,541 ± 4,210 (44,213 ± 3,916) |
| II-is | AB2 | AS | 150,618 ± 4,582 (140,074 ± 4,261) | 158,471 (147,378) | 132,635 ± 6,564 (123,351 ± 6,105) | 138,754 ± 2,885 (129,041 ± 2,683) | 136,821 ± 4,259 (127,243 ± 3,961) |
| | | SS | 88,885 ± 26,925 (82,663 ± 25,040) | 66,768 (62,095) | 85,382 ± 16,411 (79,405 ± 15,262) | 70,965 ± 4,789 (65,997 ± 4,454) | 76,556 ± 6,632 (71,197 ± 6,167) |
| | AB7 | AS | 109,826 ± 4,449 (102,138 ± 4,138) | 54,651 ± 7,680 (50,826 ± 7,142) | 110,825 ± 8,221 (103,067 ± 7,645) | 95,052 ± 3,873 (88,398 ± 3,602) | 103,746 ± 3,556 (96,484 ± 3,307) |
| | | SS | 55,540 ± 4,129 (51,652 ± 3,840) | - | 57,479 ± 5,862 (53,456 ± 5,451) | 49,167 ± 3,658 (45,725 ± 3,402) | 54,719 ± 3,688 (50,888 ± 3,430) |

| | | | | | | | |
|---|---|---|---|---|---|---|---|
| | M16 | AS | 117,058 ± 2,832 (108,864 ± 2,634) | 62,887 ± 827 (58,485 ± 769) | 108,243 ± 2,285 (100,666 ± 11,425) | 97,272 ± 8,356 (90,463 ± 7,771) | 107,303 ± 3,481 (99,791 ± 3,237) |
| | | SS | 55,689 ± 17,096 (51,791 ± 15,900) | 74,992 (69,742) | 75,836 ± 15,812 (70,528 ± 14,705) | 90,443 ± 3,183 (84,112 ± 2,960) | 75,776 ± 12,585 (70,472 ± 11,704) |
| II-ni | AB2 | AS | 134,313 ± 18,124 (124,912 ± 16,855) | 76,356 (71,011) | 142,178 ± 3,698 (132,226 ± 12,739) | 113,555 ± 3,135 (105,606 ± 2,915) | 124,586 ± 2,642 (115,865 ± 2,457) |
| | | SS | 158,933 (147,808) | - | 78,060 ± 22,575 (72,596 ± 20,995) | 65,493 ± 5,453 (60,909 ± 5,071) | 71,043 ± 6,070 (66,070 ± 5,645) |
| | AB7 | AS | 107,195 ± 6,397 (99,691 ± 5,950) | 68,166 (63,395) | 127,407 ± 14,624 (118,489 ± 13,600) | 80,975 ± 8,544 (75,306 ± 7,946) | 108,497 ± 15,681 (100,902 ±14,583) |
| | | SS | 34,529 ± 2,313 (32,112 ± 2,151) | 37,320 (34,708) | 55,050 ± 1,687 (51,197 ± 1,569) | 74,271 ± 30,748 (69,072 ± 28,596) | 78,389 ± 28,654 (72,902 ± 26,648) |
| | M16 | AS | 132, 334 ± 16,181 (123,071 ± 15,048) | 63,233 (58,807) | 147,279 ± 5,426 (136,970 ± 5,046) | 131,328 ± 6,627 (122,135 ± 6,163) | 140,641 ± 5,337 (130,796 ± 4,963) |
| | | SS | 31,209 (29,024) | 48,777 (45,363) | 84,956 ± 7,781 (79,009 ± 7,237) | 77,577 ± 16,274 (72,147 ± 15,135) | 83,259 ± 4,851 (77,431 ± 4,511) |
| III | AB2 | AS | 147,924 ± 9,132 (140,074 ± 4,261) | - | 150,023 ± 15,852 (139,521 ± 14,742) | 145,951 ± 6,739 (135,734 ± 6,267) | 148,328 ± 11,445 (137,945 ± 10,644) |
| | | SS | 249,328 (231,875) | - | 101,092 ± 10,065 (94,015 ± 9,360) | 82,422 ± 16,741 (76,653 ± 15,569) | 87,154 ± 1,548 (81,053 ± 1,440) |
| | AB7 | AS | 143,059 ± 17,823 (130,782 ± 16,576) | 60,085 ± 26,927 (55,879 ± 25,042) | 142,772 ± 22,265 (132,778 ± 20,706) | 98,564 ± 10,477 (91,665 ± 9,744) | 117,208 ± 14,761 (109,003 ± 13,728) |

| | | | | | | | |
|---|---|---|---|---|---|---|---|
| | | SS | 54,269 (50,470) | - | 56,503 ± 775 (52,547 ± 721) | 51,868 ± 15,681 (48,238 ± 14,583) | 52,796 ± 8,592 (49,100 ± 7,991) |
| | M16 | AS | 138,478 ± 15,646 (122,066 ± 14,551) | 80,726 ± 23,662 (75,075 ± 22,006) | 126,737 ± 7,554 (117,866 ± 7,025) | 136,958 ± 3,963 (127,371 ± 3,686) | 129,920 ± 6,195 (120,825 ± 5,761) |
| | | SS | 74,200 ± 13,394 (69,006 ± 12,457) | - | 80,604 ± 25,168 (74,962 ± 23,406) | 55,753 ± 5,973 (51,850 ± 5,555) | 77,780 ± 22,130 (72,336 ± 20,581) |
| Va/b | AB2 | AS | 166,753 ± 8,190 (155,081 ± 7,617) | 80,201 ± 11,699 (74,587 ± 10,880) | 165,654 ± 4,971 (154,058 ± 4,623) | 150,719 ± 7,462 (140,169 ± 6,939) | 158,490 ± 5,898 (147,395 ± 5,486) |
| | | SS | 71,678 (66,660) | 12,298 (11,437) | 63,876 ± 6,495 (59,405 ± 6,041) | 76,063 ± 25,752 (70,739 ± 23,950) | 67,725 ± 5,624 (62,984 ± 5,230) |
| | AB7 | AS | 127,985 ± 10,154 (119,026 ± 9,443) | 101,976 ± 32,066 (94,837 ± 29,821) | 123,017 ± 13,268 (114,406 ± 12,339) | 122,163 ± 5,284 (113,612 ± 4,915) | 123,224 ± 9,182 (114,598 ± 8,539) |
| | | SS | - | 53,334 (49,601) | 59,023 ± 13,066 (54,891 ± 12,151) | 98,873 ± 22,517 (91,952 ± 20,940) | 77,956 ± 15,833 (72,499 ± 14,724) |
| | M16 | AS | 151,451 ± 9,819 (140,850 ± 9,131) | 13,079 ± 504 (12,164 ± 469) | 164,896 ± 22,389 (153,353 ± 20,822) | 138,745 ± 8,249 (129,032 ± 7,672) | 147,653 ± 11,503 (137,317 ± 10,698) |
| | | SS | - | - | 55,914 ± 9,046 (52,000 ± 8,412) | 70,119 ± 893 (65,210 ± 831) | 57,871 ± 8,473 (53,820 ± 7,880) |
| Vc | AB2 | AS | 139,246 ± 8,631 (129,499 ± 8,027) | 56,969 ± 12,572 (52,981 ± 11,692) | 136,775 ± 10,553 (127,200 ± 9,814) | 128,880 ± 4,273 (119,858 ± 3,974) | 131,651 ± 6,311 (122,436 ± 5,869) |
| | | SS | 48,887 ± 2,496 (45,465 ± 2,321) | - | 90,867 ± 17,900 (84,506 ± 16,647) | 80,884 ± 11,685 (75,222 ± 10,867) | 88,640 ± 8,926 (82,435 ± 8,302) |

| | | | | | | | |
|---|---|---|---|---|---|---|---|
| | AB7 | AS | 117,786 ± 9,871 (109,541 ± 9,180) | 52,166 ± 26,276 (48,514 ± 24,437) | 129,878 ± 6,230 (120,786 ± 5,794) | 143,514 ± 23,352 (133,468 ± 21,718) | 132,248 ± 13,548 (122,991 ± 12,600) |
| | | SS | 84,649 ± 18,704 (78,723 ± 17,395) | - | 102,861 ± 22,263 (95,660 ± 20,705) | 88,327 ± 9,312 (82,144 ± 8,660) | 91,241 ± 15,165 (84,854 ± 14,103) |
| | M16 | AS | 127,954 ± 12,626 (118,997 ± 11,742) | 73,930 ± 10,687 (68,755 ± 9,939) | 140,264 ± 13,012 (130,446 ± 12,101) | 123,457 ± 8,479 (114,815 ± 7,886) | 126,953 ± 7,026 (118,066 ± 6,534) |
| | | SS | 94,996 ± 15,517 (88,346 ± 14,430) | 115,738 (107636) | 95,910 ± 24,071 (89,197 ± 22,386) | 88,869 ± 4,303 (82,648 ± 4,002) | 94,847 ± 8,257 (88,208 ± 7,679) |
| VI | AB2 | AS | 123,846 ± 17,263 (115,176 ± 16,055) | 55,467 ± 295 (51,585 ± 274) | 98,618 ± 19,859 (91,715 ± 18,469) | 134,501 ± 12,197 (125,086 ± 11,343) | 117,723 ± 12,160 (109,482 ± 11,309) |
| | | SS | - | - | 118,882 ± 22,081 (110,560 ± 20,535) | 36,961 ± 22,025 (34,374 ± 20,484) | 81,725 ± 22,427 (76,004 ± 20,857) |
| | AB7 | AS | 79,798 ± 3,452 (74,212 ± 3,211) | - | 65,899 ± 11,109 (61,286 ± 10,332) | 69,969 ± 6,480 (65,072 ± 6,026) | 69,715 ± 5,101 (64,835 ± 4,744) |
| | | SS | 33,245 ± 18,289 (30,918 ± 17,009) | - | 55,862 ± 1,095 (51,952 ± 1,018) | 29,850 ± 10,639 (27,760 ± 9,894) | 42,965 ± 11,940 (39,958 ± 11,104) |
| | M16 | AS | 114,201 ± 6,212 (106,207 ± 5,777) | 35,917 (33,402) | 134,737 ± 13,417 (125,306 ± 12,478) | 113,028 ± 12,160 (105,116 ± 8,921) | 124,061 ± 10,709 (115,377 ± 9,959) |
| | | SS | - | 52,193 (48,540) | 102,108 ± 5,441 (94,960 ± 5,061) | 75,839 ± 11,794 (70,531 ± 10,968) | 86,337 ± 10,752 (80,294 ± 10,000) |

**Supplementary File 1m. SAS area of asymmetric (AS) and symmetric (SS) synapses according to the post-synaptic target in each MEC layer, per individual case.** Data in parentheses are not corrected for shrinkage. Values (in $nm^2$) are expressed as mean ± standard error.

| Layer | % CV Synaptic density | % CV Proportion of AS | % CV AS SAS area | % CV Proportion of macular AS | % CV Proportion of AS on spines |
|---|---|---|---|---|---|
| **I** | 12.2 | 1.4 | 10.2 | 6.8 | 5.7 |
| **II-is** | 11.0 | 2.1 | 15.8 | 3.5 | 9.1 |
| **II-ni** | 12.9 | 3.6 | 14.2 | 5.2 | 15.6 |
| **III** | 18.7 | 2.7 | 13.2 | 11.2 | 11.0 |
| **Va/b** | 13.9 | 1.3 | 12.3 | 7.8 | 17.8 |
| **Vc** | 20.4 | 2.4 | 12 | 3.6 | 12.1 |
| **VI** | 14.8 | 2.2 | 19.4 | 5.3 | 4.4 |

**Supplementary File 1n. Coefficient of Variation of the analyzed synaptic parameters in each MEC layer between the stack of images.** AS: asymmetric synapses; CV: coefficient of variation; SAS: synaptic apposition surface.

| Region / Layer | Animal | Reference | Microscopy | No. Case/Sex/Age | AS Spine Heads (%) | AS Spine Necks (%) | AS Spiny Shafts (%) | AS Aspiny Shafts (%) | SS Spine Heads (%) | SS Spine Necks (%) | SS Spiny Shafts (%) | SS Aspiny Shafts (%) |
|---|---|---|---|---|---|---|---|---|---|---|---|---|
| **CEM I-VI** | Human | Present Study | FIB/SEM | 2M 1F (40-66) | 59.3 | 0.5 | 16.5 | 17.8 | 0.7 | 0.1 | 2.8 | 2.3 |
| **CE II** | Human | Domínguez-Álvaro et al., 2021b | FIB/SEM | 4M (40-66) | 53.8 | 0.5 | 19.3 | 18.2 | 1.2 | 0.1 | 5.1 | 1.9 |
| **CE III** | Human | Domínguez-Álvaro et al., 2021b | FIB/SEM | 4M (40-66) | 51 | 0.7 | 24.8 | 16.2 | 1.2 | 0.2 | 4.2 | 1.6 |
| **CTE II** | Human | Domínguez-Álvaro et al., 2019 | FIB/SEM | 4M 1F (36-66) | 54.7 | 0.5 | 19 | 18.4 | 0.5 | 0.1 | 4 | 2.8 |
| **BA21 III** | Human | Cano-Astorga et al., 2023 | FIB/SEM | 2M 1F (53-66) | 69.4 | 1.1 | 11.8 | 10.8 | 1.2 | 0.3 | 3.4 | 2.4 |
| **BA24 III** | Human | Cano-Astorga et al., 2023 | FIB/SEM | 2M 1F (53-66) | 68.5 | 1 | 13.2 | 10.8 | 1.3 | 0.6 | 2.2 | 2.5 |
| **vBA38 III** | Human | Cano-Astorga et al., 2023 | FIB/SEM | 2M 1F (53-66) | 73.4 | 0.6 | 10.6 | 10.2 | 1.2 | 0.3 | 2.6 | 1.1 |
| **dBA38 III** | Human | Cano-Astorga et al., 2023 | FIB/SEM | 2M 1F (53-66) | 71.3 | 0.8 | 11.9 | 10.1 | 1.1 | 0.3 | 2.6 | 1.9 |
| **CA1 (All Layers)** | Human | Montero-Crespo et al., 2020 | FIB/SEM | 3M 2F (36-65) | 77.8 | 0.2 | 5.6 | 5.8 | 0.9 | 0.2 | 6.2 | 5.8 |

| | | | | | | | | | | | | |
|---|---|---|---|---|---|---|---|---|---|---|---|---|
| **Temporal Cortex V** | Human | Yakuobi et al., 2019 | TEM | 3M 4F (20-50) | 85* | - | 15*(*) | - | - | - | - | - |
| **Temporal Cortex VI** | Human | Schmuhl-Giesen et al., 2022 | TEM | 2M 2F (25-63) | 80* | - | 20*(*) | - | - | - | - | - |
| **Somatosensory Cortex I-VI** | Mouse | Turégano-López, 2022 | FIB/SEM | 3M (8 weeks old) | 74.1 | 2 | 10.3 | 2.3 | 3.55 | - | 7.4 | 0.5 |
| **Somatosensory Cortex I-VI** | Etruscan Shrew | Alonso-Nanclares et al., 2023 | FIB/SEM | 3M (8-20 months old) | 77.9*** | - | 13.9** | - | 0.8*** | - | 7.1** | - |
| **Somatosensory Cortex I-VI** | Juvenile Rat | Santuy et al., 2018 | FIB/SEM | 3M (14 days old) | 73.9 | 1.6 | 14.7*** | - | 2 | 0.7 | 7*** | - |

**Supplementary File 1o. Data on postsynaptic targets in different species, regions and cortical layers.** Abbreviations: EC: Entorhinal Cortex; F: Female; FIB/SEM: Focused Ion Beam-Scanning Electron Microscopy; M: Male; MEC: Medial Entorhinal Cortex; TCE: Transentorhinal Cortex; TEM: Transmission Electron Microscopy. Age refers to years, except otherwise indicated. *No classification of the type of synapses (AS:SS) were performed. **Only axospinous synapses (established on dendritic spine heads and necks) and axodendritic synapses (formed on dendritic shafts) are indicated. ***No distinction between spiny and aspiny shafts were made.

| Case | Sex | Age (years) | Cause of death | Post-mortem delay (h) | Neurological diagnosis |
|---|---|---|---|---|---|
| AB2 | Female | 53 | Pulmonary Shock | 4 | No neurological alterations |
| AB7 | Male | 66 | Metastatic Bladder Carcinoma | 2.4 | No neurological alterations |
| M16 | Male | 40 | Car Accident | 3 | No neurological alterations |

**Supplementary File 1p.** Clinical and neuropsychological information from the cases analyzed.

**Legend to Supplementary File 2.**

Supplementary File 2. Spreadsheet containing the spatial extents of individual stacks of images, detailed per case and layer. Original dimensions (x, y, z), original volumes, counting frame (CF) dimensions and CF volume are provided.

**Legend to Supplementary File 3.**

Supplementary File 3. Spreadsheet containing Kruskal-Wallis and Dunn's tests p-values of the interindividual variability analysis, organized in TABS as follows:
SYNAPTIC DENSITY
INTERSINAPTIC DISTANCE
SAS AREA OF AS
SAS AREA OF SS
SAS AREA OF MACULAR AS
SAS AREA OF MACULAR SS
SAS AREA OF COMPLEX-SHAPED AS
SAS AREA OF COMPLEX-SHAPED SS
The criterion for statistical significance was considered to be met if $p<0.05$ (indicated in the spreadsheet as *) or $p<0.01$ (**).

**Legend to Supplementary File 4.**

Supplementary File 4. Spreadsheet containing the complete raw dataset organized in TABS as follows:
RAW DATA SYNAPSES: each row corresponds to each analyzed synapse and contains its characteristics, SAS area corrected for tissue shrinkage and SAS area provided by EspINA
RAW DATA DENSITIES: each row corresponds to each analyzed stack of images and contains the number of synapses per type, the volume of the counting frame, the volume of the counting frame corrected for tissue shrinkage and artifacts, and the calculated synaptic density per type and total
SIZE: mean SAS area, SE, and number of synapses calculated per stack of images, per synaptic type and layer
SHAPE: number of synapses, mean SAS area and SE per synaptic shape, calculated per stack of images, synaptic type and layer
POSTSYNAPTIC: number of synapses, mean SAS area and SE per postsynaptic target, calculated per stack of images, synaptic type and layer
